# Presynaptic Gα_o_ (GOA-1) signals to depress command neuron excitability and allow stretch-dependent modulation of egg laying in *Caenorhabditis elegans*

**DOI:** 10.1101/701664

**Authors:** Bhavya Ravi, Jian Zhao, Sana Chaudhry, Mattingly Bartole, Richard J. Kopchock, Christian Guijarro, Lijun Kang, Kevin M. Collins

## Abstract

Egg laying in the nematode worm *Caenorhabditis elegans* is a two-state behavior modulated by internal and external sensory input. We have previously shown that homeostatic feedback of embryo accumulation in the uterus regulates bursting activity of the serotonergic HSN command neurons that sustains the egg-laying active state. How sensory feedback of egg release signals to terminate the egg-laying active state is less understood. We find that Gα_o_, a conserved Pertussis Toxin-sensitive G protein, signals within HSN to inhibit egg-laying circuit activity and prevent entry into the active state. Gα_o_ signaling hyperpolarizes HSN, reducing HSN Ca^2+^ activity and input onto the postsynaptic vulval muscles. Loss of inhibitory Gα_o_ signaling uncouples presynaptic HSN activity from a postsynaptic, stretch-dependent homeostat, causing precocious entry into the egg-laying active state when only a few eggs are present in the uterus. Feedback of vulval opening and egg release activates the uv1 neuroendocrine cells which release NLP-7 neuropeptides which signal to inhibit egg laying through Gα_o_-independent mechanisms in the HSNs and Gα_o_-dependent mechanisms in cells other than the HSNs. Thus, neuropeptide and inhibitory Gα_o_ signaling maintains a bi-stable state of electrical excitability that dynamically controls circuit activity in response to both external and internal sensory input to drive a two-state behavior output.

## Introduction

A major goal of neuroscience is to understand how sensory signals control neural circuit activity and changes in animal behavior. Such sensory feedback informs when a behavior should begin, how long it should continue, and when it should end. Extensive evidence has shown that neuromodulators like serotonin and neuropeptides signal through G protein coupled receptors to remodel neural circuit activity and drive behavior state transitions (Jiang et al., 2001; Goulding et al., 2008). Yet, there is no neural circuit in any organism for which we know precisely how each signaling event contributes to sensory modulation of a behavior. Small neural circuits typically found in invertebrates combine anatomical simplicity with genetic and experimental accessibility, allowing for a complete understanding of the molecular and physiological basis for a behavioral output (Marder, 2012).

The *C. elegans* female egg-laying behavior circuit is ideally suited to study how sensory signals modulate circuit functions that underlie decision-making. As shown in Figure 1A, the circuit is comprised of two Hermaphrodite Specific Neurons (HSNs) that synapse onto the egg-laying vulval muscles (White et al., 1986; Shen et al., 2004; Li et al., 2013). During ~2 minute active states, rhythmic HSN Ca^2+^ activity releases serotonin and neuropeptides that signal to promote the excitability of the muscles, driving ejection of ~4-6 eggs in sequence from the uterus into the environment (Waggoner et al., 1998; Shyn et al., 2003; Zhang et al., 2008; Collins et al., 2016; Brewer et al., 2019). External and internal sensory input regulate the onset of egg laying (Horvitz et al., 1982; Trent, 1982; Sawin, 1996; Aprison and Ruvinsky, 2014; Ravi et al., 2018a), and genetic studies have identified neuropeptides, receptors, and two antagonistic heterotrimeric G proteins that signal to regulate HSN activity, neurotransmitter release, and egg laying (Schafer, 2006; Koelle, 2016). Gα_q_ signals through the conserved PLCβ and Trio RhoGEF effector pathways to promote neurotransmitter release and egg laying (Brundage et al., 1996; Lackner et al., 1999; Miller et al., 1999; Bastiani et al., 2003; McMullan et al., 2006; Williams et al., 2007; McMullan and Nurrish, 2011). Because phorbol ester DAG-mimetics such as PMA rescue synaptic transmission defects of Gα_q_ signaling mutants (Lackner et al., 1999; Williams et al., 2007), DAG production and recruitment of UNC-13 and/or Protein Kinase C effectors is thought to mediate the Gα_q_ dependent modulation of synaptic transmission (Yawo, 1999; Wierda et al., 2007; Lou et al., 2008). Gα_q_ signaling is opposed by Gα_o_ which signals to inhibit neurotransmitter release (Koelle and Horvitz, 1996; Miller et al., 1999; Nurrish et al., 1999), vulval muscle activity (Shyn et al., 2003), and egg laying (Mendel et al., 1995; Segalat et al., 1995). How Gα_o_ signaling antagonizes Gα_q_, Rho, and DAG signaling is not clear. Even though Gα_o_ mediates signaling by numerous neurotransmitters in diverse animals including mammals (Jiang et al., 2001), direct effectors for Gα_o_ have not yet been identified. In *C. elegans*, Gα_o_ mutants resemble animals lacking DAG Kinase; both mutants show increased UNC-13 localization to synapses and hyperactive egg-laying behavior defects, resembling treatment with phorbol esters (Miller et al., 1999; Nurrish et al., 1999; Jose and Koelle, 2005). Because synaptic transmission could be modulated by upstream changes in cell and circuit electrical excitability and/or by downstream effects on synaptic vesicle fusion, what is needed are direct measurements of how discrete changes in G protein and effector signaling affect cell and circuit activity and their consequences on animal behavior.

**Figure 1.**
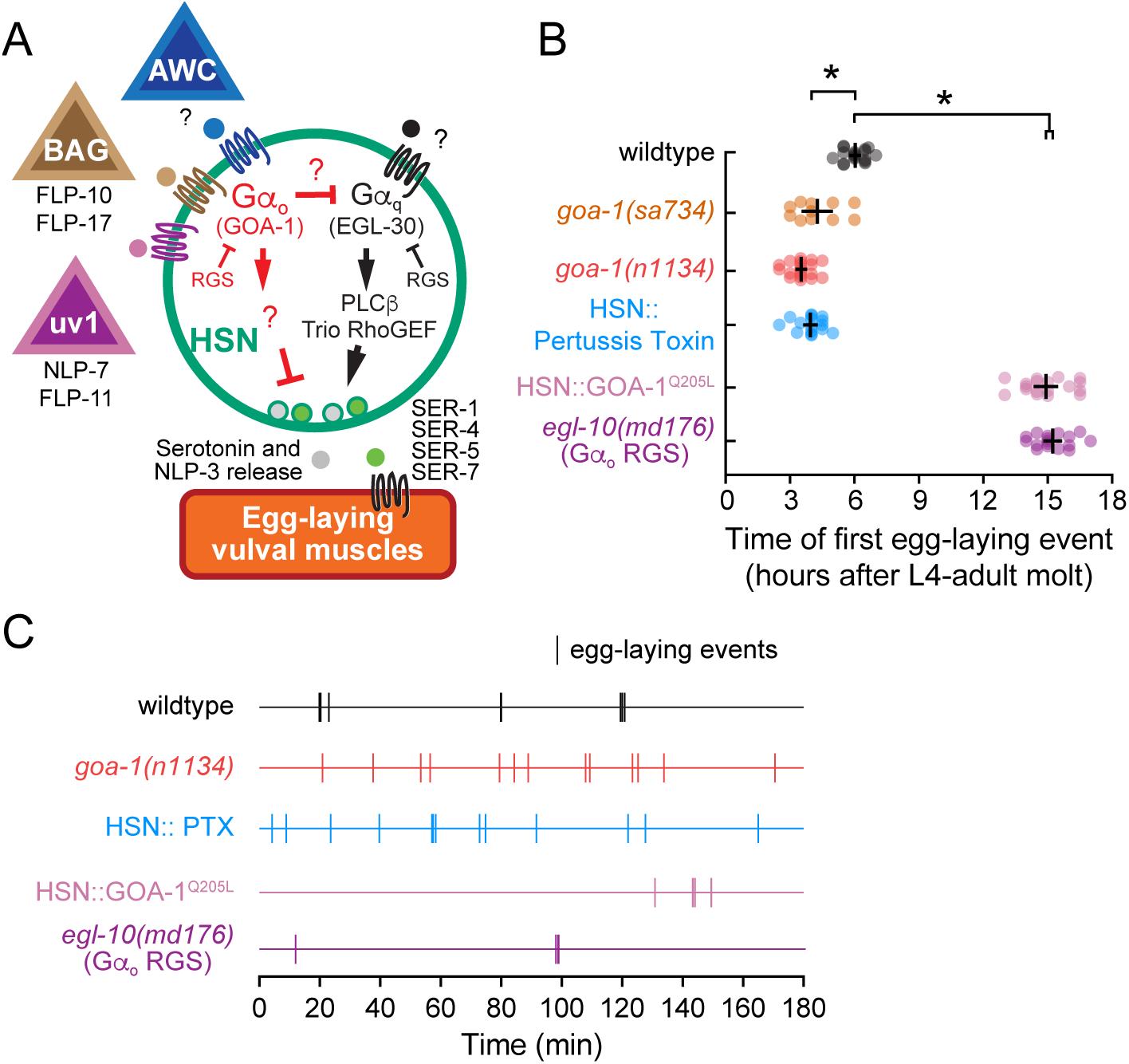
Gα_o_ signaling maintains the inactive egg-laying behavior state. (**A**) Cartoon of how identified and unidentified neuropeptides released from AWC (Fenk and de Bono, 2015), BAG (Ringstad and Horvitz, 2008), uv1 (Banerjee et al., 2017), and other sensory cells bind to G-protein coupled receptors expressed on HSN command neurons (green) which signal via Gα_o_ (red) or Gα_q_ (black) effector pathways to modulate serotonin and NLP-3 neuropeptide release (Tanis et al., 2008; Brewer et al., 2019). The egg-laying vulval muscles (orange) express receptors for serotonin (Carnell et al., 2005; Dempsey et al., 2005; Hobson et al., 2006; Fernandez et al., 2020) and possibly NLP-3 which signal to promote vulval muscle excitability and egg laying (**B**) Scatter plots of the first egg-laying event in wildtype (grey), null *goa-1(sa734)* mutants (orange), hypomorphic loss-of-function *goa-1(n1134)* mutants (red), *egl-10(md176)* null mutants (purple), and transgenic animals expressing Pertussis Toxin (blue) or GOA-1^Q205L^ in the HSNs (pink). Error bars show 95% confidence intervals for the mean from ≥10 animals. Asterisks indicate p≤0.0001 (One-way ANOVA with Bonferroni correction for multiple comparisons). (**C**) Representative raster plots showing temporal pattern of egg laying during three hours in wild-type (black), hypomorphic loss-of-function *goa-1(n1134)* mutant (red), and *egl-10(md176)* null mutant animals (purple), along with transgenic animals expressing Pertussis Toxin (blue) or GOA-1^Q205L^ in the HSNs (pink). Vertical lines indicate single egg-laying events.

Here we explore how Gα_o_ inhibits *C. elegans* egg-laying circuit activity and behavior. Our data reveal that Gα_o_ signaling reduces the electrical excitability of the HSN command neurons to promote the inactive state while eggs are produced. Feedback of egg accumulation in the uterus then switches the circuit into active states where rhythmic bursting drives sequential egg-laying events. Thus, modulation by inhibitory Gα_o_ signaling allows for proper induction of motor behavior circuit upon the alignment of optimal external and internal sensory conditions.

## Results

### Reduced inhibitory Gα_o_ signaling leads to premature egg laying and decreases the duration of egg-laying inactive states

To better understand how inhibitory Gα_o_ signaling contributes to egg-laying behavior state transitions, we used mutants and transgenes to manipulate Gα_o_ signaling in specific cells of the egg-laying circuit and analyzed the consequences on cell Ca^2+^ activity and behavior. We found that Gα_o_ signaling inhibits the onset of egg laying. We performed a ‘time to first egg lay’ assay in wild-type animals and in mutants with excessive or reduced Gα_o_ signaling. As previously shown, wild-type animals release their first embryo ~6-7 hours after becoming adults (McMullen et al., 2012; Ravi et al., 2018a). Animals bearing Gα_o_ loss-of-function or null mutations laid their eggs much earlier, 3-4 hours after becoming adults (Figure 1B). *goa-1(n1134)*, a hypomorphic mutant predicted to lack the conserved N-terminal myristoylation and palmitoylation sequence, and *goa-1(sa734)*, an early stop mutant predicted to be a molecular null (Segalat et al., 1995; Robatzek and Thomas, 2000), showed similarly precocious onset of egg laying (Figure 1B). This phenotype was shared in transgenic animals where Gα_o_ function was inhibited just in HSNs through the cell-specific expression of Pertussis Toxin (Tanis et al., 2008). Because the timing of this first egg-laying event requires serotonin along with HSN function and activity (Ravi et al., 2018a), these results suggest that Gα_o_ normally signals in HSN to inhibit neurotransmitter release and thereby delay the first egg-laying active state (Figure 1B). To test the effects of increased Gα_o_ signaling, we analyzed the behavior of *egl-10(md176)* mutants which lack the major RGS protein that terminates Gα_o_ signaling by promoting Gα_o_ GTP hydrolysis (Koelle and Horvitz, 1996). *egl-10(md176)* mutants showed a strong and significant delay in the onset of egg laying, laying their first egg ~15 hours after reaching adulthood (Figure 1B). This delay of egg-laying phenotype was shared in transgenic animals expressing the constitutively active Gα_o_ (Q205L) mutant specifically in the HSNs (Tanis et al., 2008) and resembled animals developmentally lacking HSNs (Ravi et al., 2018a), consistent with Gα_o_ signaling in HSN to inhibit neurotransmitter release that normally drives the onset of egg laying.

To understand how Gα_o_ signaling controls the normal two-state pattern of egg laying, we performed long-term recordings of adults as they transitioned into and out of the active states in which clusters of several eggs are typically laid. We defined intra-cluster intervals as the time elapsed between consecutive egg-laying events within a single active state (e.g. intervals < 4 minutes) and inter-cluster intervals as the time elapsed between different active states (e.g. intervals > 4 minutes), as described (Waggoner et al., 1998; Collins and Koelle, 2013). As expected, wild-type animals displayed a two-state pattern of egg laying with multiple egg-laying events clustered within brief, ~2-minute active states about every 20-30 minutes (Figure 1C and Table 1). Animals bearing the *goa-1(n1134)* hypomorphic mutation entered active states 2-3-fold more frequently, often laying single eggs during active states separated by only ~12-13 minutes (Figure 1C and Table 1), as previously shown (Waggoner et al., 2000). We found the duration of inactive states in animals expressing Pertussis Toxin in the HSN neurons was indistinguishable from the *goa-1(n1134)* mutant, indicating that Gα_o_ signals in HSN to reduce the probability of entering the active state (Figure 1C and Table 1). Loss of inhibitory Gα_o_ signaling led to active states in which 1-2 embryos were laid almost immediately after they were positioned in the uterus next to the vulval opening. As a result, successive egg-laying events were rate-limited by egg production, explaining why the average intra-cluster intervals were typically double seen in wild-type animals (Figure 1C and Table 1). Conversely, *egl-10(md176)* mutant animals or transgenic animals expressing the Gα_o_(Q205L) gain-of-function mutation in the HSNs showed prolonged inactive periods of 258 and 67 min, respectively (Figure 1C and Table 1), resembling animals lacking HSNs altogether (Waggoner et al., 2000). Interestingly, animals without HSNs or with too much inhibitory Gα_o_ signaling still lay multiple eggs within discrete active states (Figure 1C and Table 1), consistent with our results showing that a stretch-dependent homeostat can maintain the active state even when neurotransmitter release from the HSN is inhibited (Collins et al., 2016; Ravi et al., 2018a). These results show that Gα_o_ signaling does not modulate patterns of egg laying within active states. Instead, Gα_o_ acts in HSN to depress entry into the egg-laying active state. Long-term behavior recordings were used to extract features of egg-laying active and inactive behavior states for the indicated genotypes, as described (Waggoner et al., 1998). ‘*’ indicates significant differences compared to wild-type animals (p<0.0001, Kruskal-Wallis test with Dunn’s correction for multiple comparisons), ‘ǂ’ indicates significant differences compared to wildtype (p<0.0001, One-way ANOVA with Bonferroni correction), and ‘#’ indicates that this result was previously reported (Tanis et al., 2008).

**Table 1.**
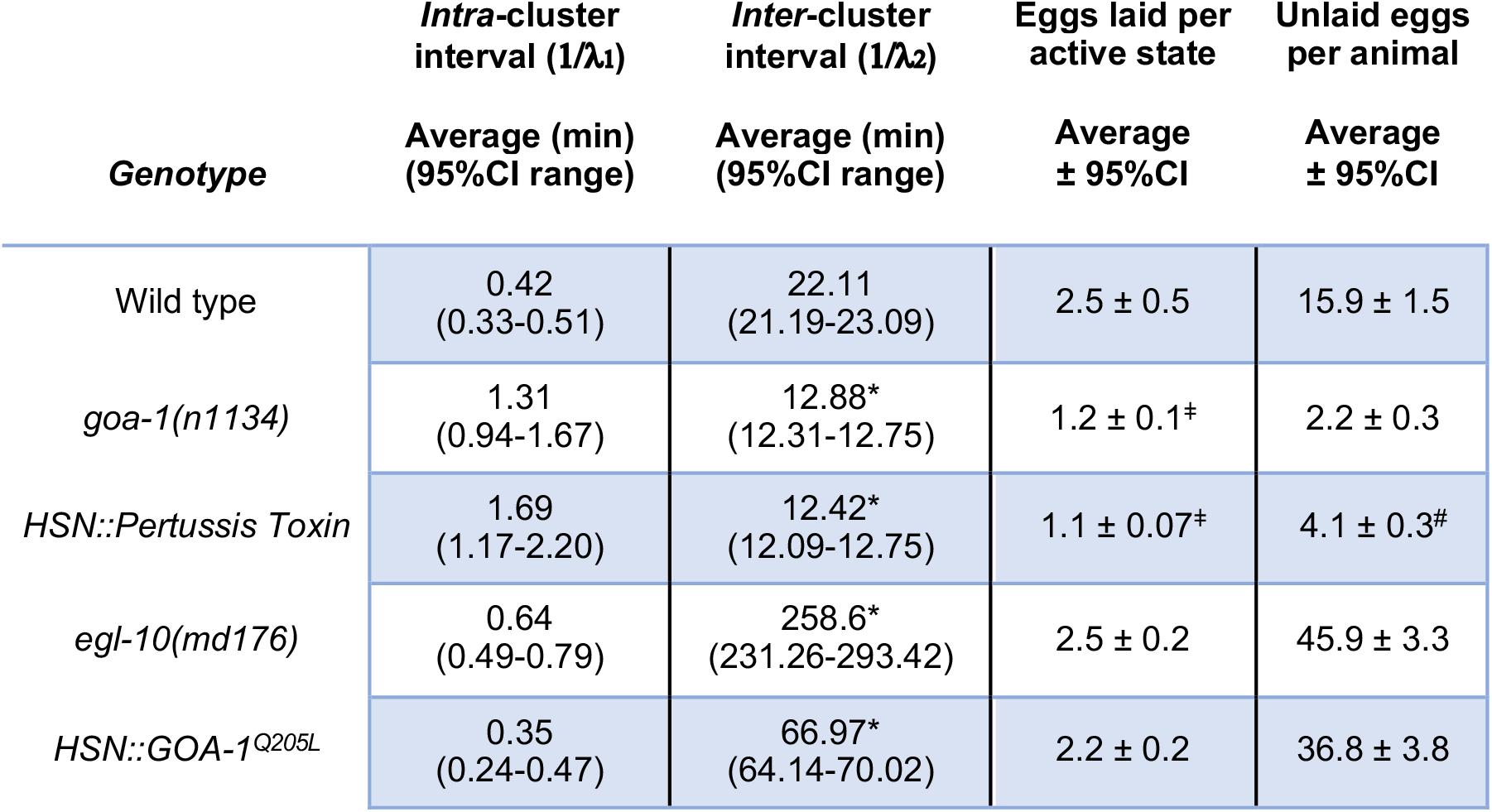
Egg-laying behavior measurements in animals with altered Gα_o_ signaling.

### Gα_o_ signaling inhibits HSN Ca^2+^ activity to promote the inactive behavior state

Previous work has shown that Gα_o_ signals presnaptically to depress neurotransmitter release. Whether Gα_o_ signals to inhibit HSN electrical excitability that might be evident in changes in cell Ca^2+^ activity, or instead signals downstream of Ca^2+^ to regulate steps in UNC-13-dependent docking, priming, and vesicle fusion, has not been tested directly. To address how Gα_o_ signals to inhibit HSN neurotransmitter release, we performed ratiometric Ca^2+^ imaging in our panel of Gα_o_ signaling mutants as they entered spontaneous egg-laying active states. Freely behaving animals bearing the *goa-1(n1134)* hypomorphic or *goa-1(sa734)* null mutations showed a clear change in HSN Ca^2+^ activity from two-state burst firing to more tonic firing (Figure 2A, Videos 1-3). Complete loss of inhibitory Gα_o_ signaling caused a significant increase in the frequency of HSN Ca^2+^ transients (Figure 2B and 2C). We were surprised that the *goa-1(n1134)* mutants, which show strongly hyperactive egg-laying behavior indistinguishable from that of *goa-1(sa734)* null mutants, showed only a modest and statistically insignificant increase in HSN Ca^2+^ activity compared to wild-type (Figure 2C). However, our results with *goa-1(n1134)* match previously findings (Shyn et al., 2003). The *goa-1(n1134)* hypomorphic mutant is expected to have residual Gα_o_ signaling activity in that its major defect is the absence of a proper membrane anchor sequence (Mumby et al., 1990). These results show that Gα_o_ signaling depresses HSN presynaptic activity, but that the hyperactive egg-laying phenotypes observed in *goa-1(n1134)* mutants are separable from an increase in presynaptic HSN Ca^2+^ activity.

**Figure 2.**
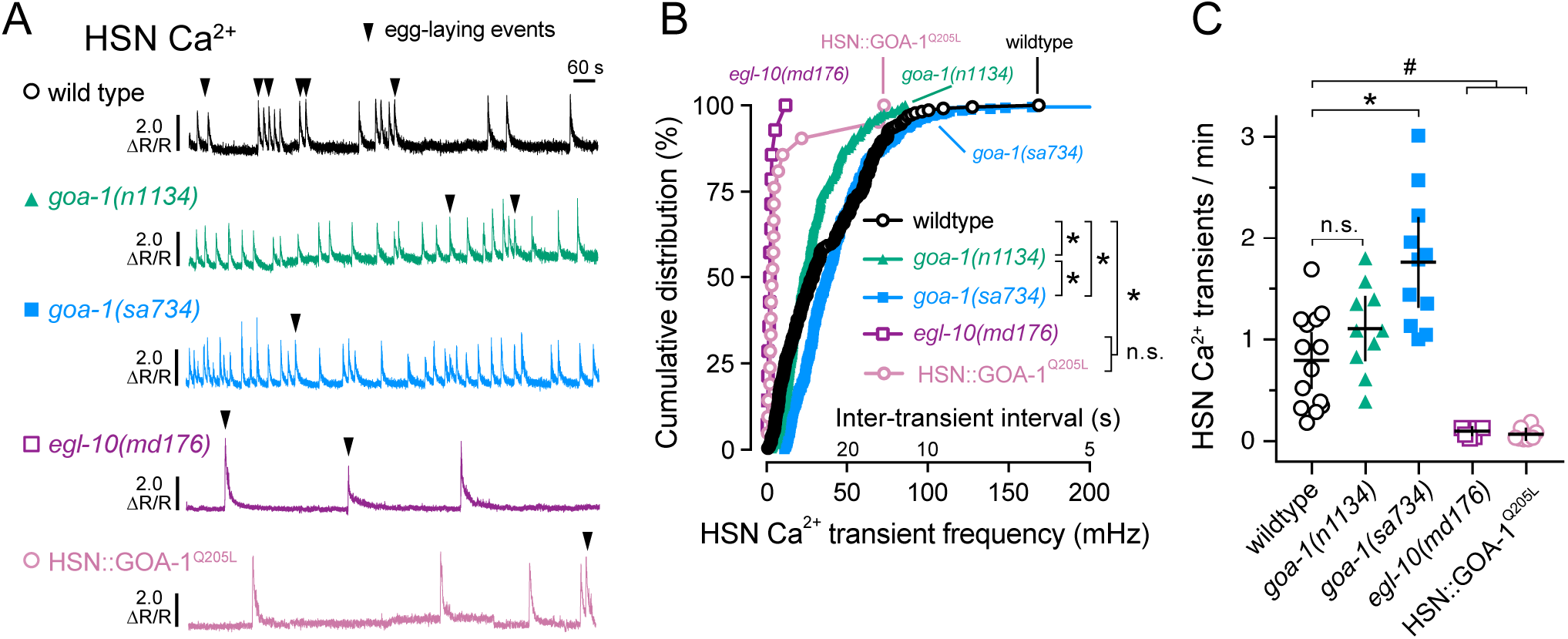
Gα_o_ signaling inhibits HSN neuron Ca^2+^ activity and burst firing. (**A**) Representative GCaMP5:mCherry (ΔR/R) ratio traces showing HSN Ca^2+^ activity in freely behaving wild-type (black), *goa-1(n1134)* loss-of-function mutant (green), *goa-1(sa734)* null mutant (blue), and *egl-10(md176)* null (purple) mutant animals, along with transgenic animals expressing the activated GOA-1(Q205L) in the HSN neurons (pink) during an egg-laying active state. Arrowheads indicate egg-laying events. (**B**) Cumulative distribution plots of instantaneous Ca^2+^ transient peak frequencies (and inter-transient intervals) in wild-type (black open circles), *goa-1(n1134)* (green filled triangles), *goa-1(sa734)* (blue squares), *egl-10(md176)* mutants (purple open squares) along with transgenic animals expressing the activated GOA-1(Q205L) in the HSN neurons (pink open circles). Asterisks indicate p<0.0001 (Kruskal-Wallis test with Dunn’s correction for multiple comparisons). (**C**) Scatter plots show average Ca^2+^ transient frequency (per min). Error bars indicate 95% confidence intervals for the mean. Asterisk indicates p<0.0001; pound (#) indicates p≤0.0079; n.s. indicates p>0.05 (One-way ANOVA with Bonferroni correction for multiple comparisons).

We next tested how increased inhibitory Gα_o_ signaling affects HSN activity. Both *egl-10(md176)* mutants and transgenic animals expressing the activated GOA-1(Q205L) in HSNs showed a significant and dramatic reduction in the frequency of HSN Ca^2+^ transients, with single HSN Ca^2+^ transients occuring several minutes apart (Figure 2A and 2B). The rare egg-laying events seen in animals with increased Gα_o_ signaling mostly followed single HSN Ca^2+^ transients, not multi-transient bursts typically seen in wild-type animals (Figure 2A and 2C). In one *egl-10(md176)* animal, we observed an egg-laying event that was not accompanied by an HSN Ca^2+^ transient. This suggests that elevated Gα_o_ signaling is sufficient to silence the HSNs, and that, in this case, egg laying becomes HSN-independent. In support of this model, previous work has shown that *egl-10(md176)* mutants respond to serotonin but are resistant to the serotonin reuptake inhibitor imipramine (Trent et al., 1983). Alternatively (or additionally) Gα_o_ signaling may function to depress coordinated activity between the gap-junctioned, contralateral HSNs, whose Ca^2+^ activity we were unable to observe simultaneously because our confocal imaging conditions only captures one HSN at a time.

To determine how disruption of inhibitory Gα_o_ signaling in HSN affects its activity, we recorded HSN Ca^2+^ transients in transgenic animals expressing Pertussis Toxin in the HSNs. Gα_o_-silenced HSNs showed a dramatic increase in the frequency of HSN Ca^2+^ transients, leading to a nearly constitutive, tonic Ca^2+^ activity like that observed in *goa-1(sa734)* null mutants (Figure 3A, 3B, and 3C; Video 4). While control animals showed an average HSN Ca^2+^ transient frequency of about ~0.4 transients per minute, animals expressing Pertussis Toxin in HSN showed an average 1.9 transients per minute, a significant increase (Figure 3C). These results suggest that even under normal growth conditions, unidentified neuromodulators signal through Gα_o_-coupled receptors on HSN to inhibit Ca^2+^ activity. We hypothesize that the observed changes in presynaptic UNC-13 localization (Nurrish et al., 1999) and serotonin biosynthesis (Tanis et al. 2008) when inhibitory Gα_o_ signaling is reduced are consequences, not causes, of altered presynaptic activity.

**Figure 3.**
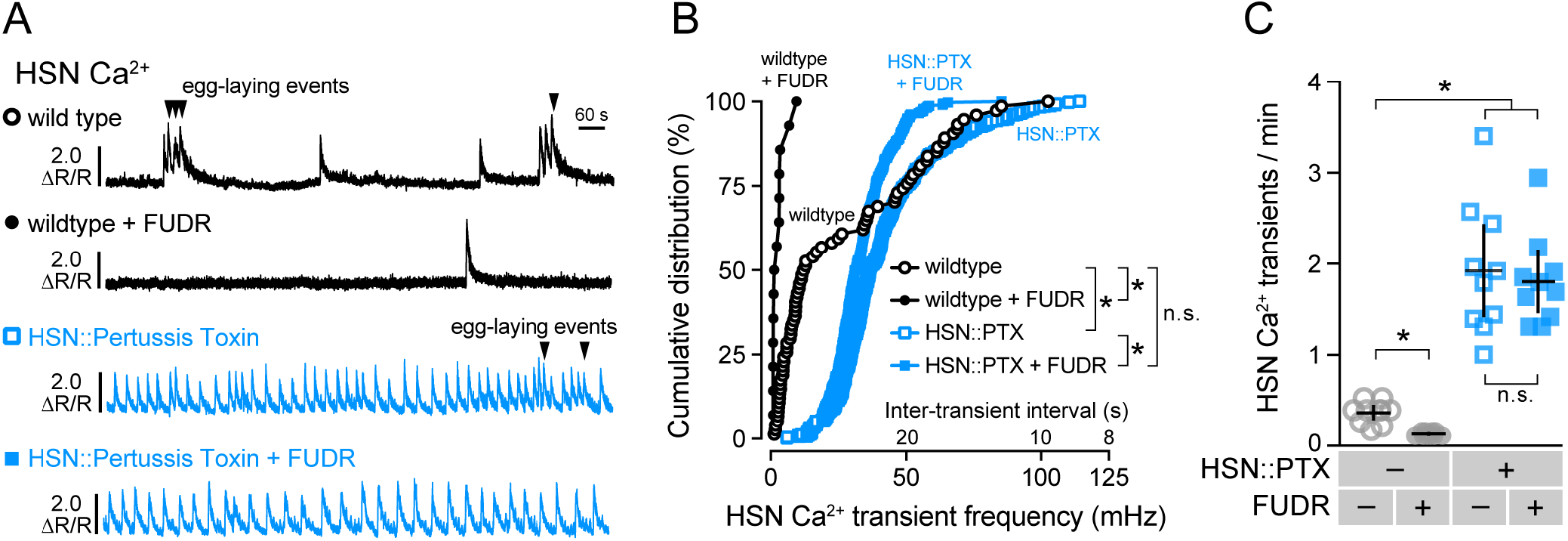
Inhibitory Gα_o_ signaling in HSN is required for two-state Ca^2+^ activity and facilitates modulation by feedback of egg accumulation. **(A)** Representative GCaMP5:mCherry (ΔR/R) ratio traces showing HSN Ca^2+^ activity in untreated (fertile) wild-type animals (black open circles), FUDR-sterilized wild-type animals (black filled circles), untreated (fertile) animals expressing Pertussis Toxin (PTX) in the HSN neurons (blue open squares), and in FUDR-sterilized transgenic animals expressing PTX in the HSNs (blue filled squares). Arrowheads indicate egg-laying events. (**B**) Cumulative distribution plots of instantaneous Ca^2+^ transient peak frequencies (and inter-transient intervals) in untreated and FUDR-treated animals. Asterisks indicate p<0.0001 (Kruskal-Wallis test with Dunn’s test for multiple comparisons). (**C**) Scatter plots show average Ca^2+^ transient frequency (per min). Error bars indicate 95% confidence intervals for the mean. Asterisks indicate p<0.0001 (One-way ANOVA with Bonferroni’s test for multiple comparisons). Data from 10 animals were used for each strain for analysis.

We have previously shown that burst Ca^2+^ activity in the command HSN neurons is initiated and sustained by a stretch-dependent homeostat. In chemically or genetically sterilized animals, burst Ca^2+^ activity in HSN is largely eliminated (Ravi et al., 2018a). As such, we were surprised to observe high frequency Ca^2+^ transients in animals with reduced Gα_o_ signaling because these animals typically retain few (1 to 3) eggs in the uterus at steady state, conditions that normally eliminate HSN burst firing. We hypothesized that the stretch-dependent homeostat was not required to promote HSN Ca^2+^ activity in Gα_o_ signaling mutants. To test this, we chemically sterilized transgenic animals expressing Pertussis Toxin in the HSNs with Floxuridine (FUDR), a blocker of embryogenesis (Mitchell et al., 1979), and recorded HSN Ca^2+^ activity. Wild-type animals treated with FUDR showed a dramatic decrease in the frequency of HSN Ca^2+^ activity and an elimination of burst firing (Figures 3A-C), as we have previously shown (Ravi et al., 2018a). Sterilized transgenic animals expressing Pertussis Toxin in the HSNs showed only a slight reduction in HSN Ca^2+^ frequency (Figure 3A and 3B). Both fertile and sterile Pertussis Toxin-expressing animals had dramatically and significantly increased HSN Ca^2+^ transient frequency (~1.9 / min), indicating their HSNs no longer responded to the retrograde signals of egg accumulation arising from the stretch-homeostat. One explanation for this result could be that in fertile wild-type animals, feedback of egg accumulation elevates HSN excitability above a firing threshold that overcomes endogenous inhibitory Gα_o_ signaling. Together, these results support a model where Gα_o_ signals in HSN to depress cell electrical excitability which allows for the proper two-state pattern of HSN Ca^2+^ activity that responds to homeostatic feedback of egg accumulation.

### Presynaptic Gα_o_ signaling inhibits postsynaptic vulval muscle activity

To test how changes in inhibitory Gα_o_ signaling affect the postsynaptic vulval muscles, we recorded Ca^2+^ activity in the vulval muscles of *goa-1(n1134)* mutant and Pertussis Toxin expressing transgenic animals. Both wild-type and *goa-1(n1134)* mutants showed increased vulval muscle Ca^2+^ transients as the animals entered the egg-laying active state (Figure 4A and 4B). *goa-1(n1134)* mutants also showed elevated muscle activity compared to wild-type control animals during inactive states when no eggs were laid (Figure 4A and 4B), as previously shown (Shyn et al., 2003). Surprisingly, Ca^2+^ activity during egg-laying active states was not significantly different in *goa-1(n1134)* mutant animals (Figure 4A and 4B; compare Videos 5 and 6), a result consistent with Gα_o_ signaling primarily to prolong the inactive state.

**Figure 4.**
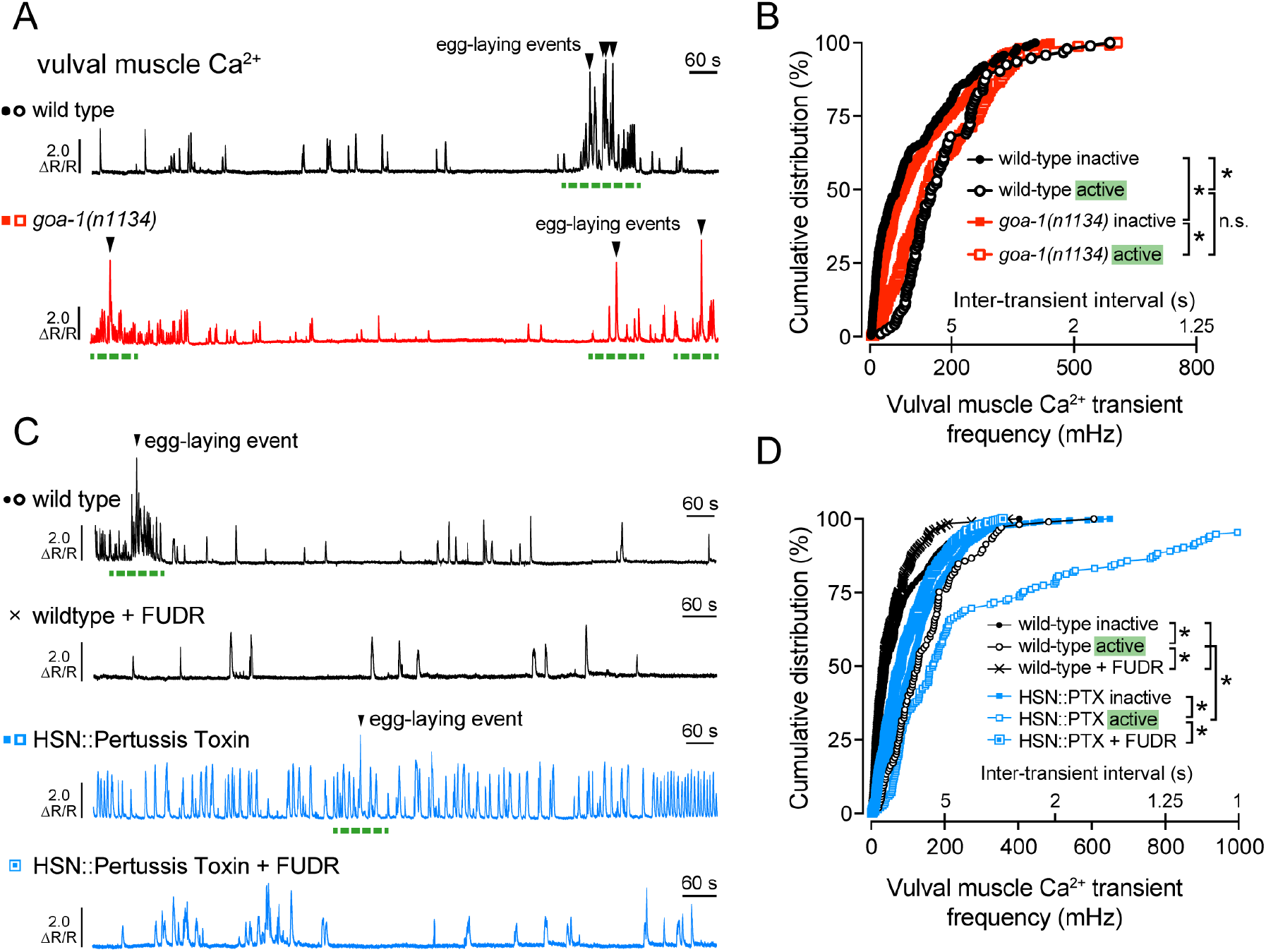
Gα_o_ signals in HSN to reduce excitatory modulation of the vulval muscles. **(A)** Representative GCaMP5:mCherry (ΔR/R) ratio traces showing vulval muscle Ca^2+^ activity in wild-type (black) and *goa-1(n1134)* loss-of-function mutant animals (red). Egg-laying events are indicated by arrowheads, and egg-laying active states are indicated by dashed green lines. **(B)** Cumulative distribution plots of instantaneous vulval muscle Ca^2+^ transient peak frequencies (and inter-transient intervals) in wild-type (black circles) and *goa-1(n1134)* mutant animals (red squares) in the inactive and active egg-laying states (filled and open, respectively). **(C)** Representative GCaMP5:mCherry (ΔR/R) ratio traces showing vulval muscle Ca^2+^ activity in untreated (circles) and FUDR-treated (cross) wild-type animals (black) along with untreated (filled) or FUDR-treated (open) transgenic animals expressing Pertussis Toxin in the HSNs (blue squares). Egg-laying events are indicated by arrowheads, and egg-laying active states are indicated by green dashed lines. **(D)** Cumulative distribution plots of instantaneous vulval muscle Ca^2+^ transient peak frequencies (and inter-transient intervals) in wildtype and in transgenic animals expressing Pertussis Toxin in the HSN neurons, with or without FUDR treatment (black cross or blue square), during the inactive and active egg-laying states (filled and open, respectively). Asterisks indicate p<0.0001 (Kruskal-Wallis test with Dunn’s test for multiple comparisons).

These results suggest Gα_o_ signals to inhibit circuit activity during the egg-laying inactive state and/or that loss of Gα_o_ elevates circuit activity that prolongs the active state beyond the brief ~2-3 minute window during which eggs are typically laid (Waggoner et al., 1998). Recordings of vulval muscle Ca^2+^ activity in transgenic animals expressing Pertussis Toxin in the presynaptic HSNs show a dramatic and significant increase in vulval muscle Ca^2+^ transient frequency during both active and inactive states (Figure 4C and 4D; Video 7), confirming Gα_o_ is required in HSN to inhibit neurotransmitter release.

We have previously shown that egg accumulation promotes vulval muscle excitability during the active state while sterilization reduces it to that seen during the inactive state (Collins et al., 2016; Ravi et al., 2018a). Because FUDR-sterilization failed to reduce HSN Ca^2+^ activity in Pertussis Toxin expressing animals (Figure 3C), we predicted that vulval muscle Ca^2+^ activity would remain similarly elevated in these animals after sterilization. To our surprise, FUDR treatment significantly reduced, but did not completely eliminate, the elevated vulval muscle Ca^2+^ activity in animals expressing Pertussis Toxin in HSN (Figure 4C and 4D). This result suggests egg accumulation and/or germline activity is still required for full vulval muscle activity even when HSN Ca^2+^ activity is dramatically increased. However, because these animals lay eggs almost as soon as they are made, the degree of uterine stretch necessary to induce the active state must be markedly reduced. This supports a model where both the stretch-dependent homeostat and Gα_o_ signaling dynamically interact to regulate egg-laying behavior states.

### Gα_o_ signaling modulates the HSN resting membrane potential

Reduction of inhibitory Gα_o_ signaling strongly increased HSN Ca^2+^ activity and burst firing, prompting us to investigate whether Gα_o_ signals to modulate HSN electrical excitability. We recorded the resting membrane potential of the HSN neurons in animals with altered Gα_o_ signaling using the whole-cell patch clamp technique (Figure 5A), as described (Yue et al., 2018). Hypomorphic *goa-1(n1134)* loss-of-function mutants displayed a trend towards more depolarized resting potentials (−17.9 mV) compared to HSNs from recorded wild-type animals (−21.1 mV), but this difference was not statistically significant (Figure 5B). In contrast, the resting membrane potential of HSNs in *egl-10(md176)* Gα_o_ RGS protein mutant animals with a global increase in Gα_o_ signaling (Koelle and Horvitz, 1996) was significantly hyperpolarized (−40.8 mV) compared to wild-type control animals. This hyperpolarization of HSNs in *egl-10(md176)* mutants explains the reduced frequency of HSN Ca^2+^ transients and their strong defects in egg-laying behavior. Transgenic animals expressing Pertussis Toxin specifically in the HSNs had significantly depolarized HSNs (−14.75 mV) compared to the wild-type parental strain (−21.8 mV). These results show that Gα_o_ signals in the HSNs to promote membrane polarization, reducing cell electrical excitability, Ca^2+^ activity, and neurotransmitter release.

**Figure 5.**
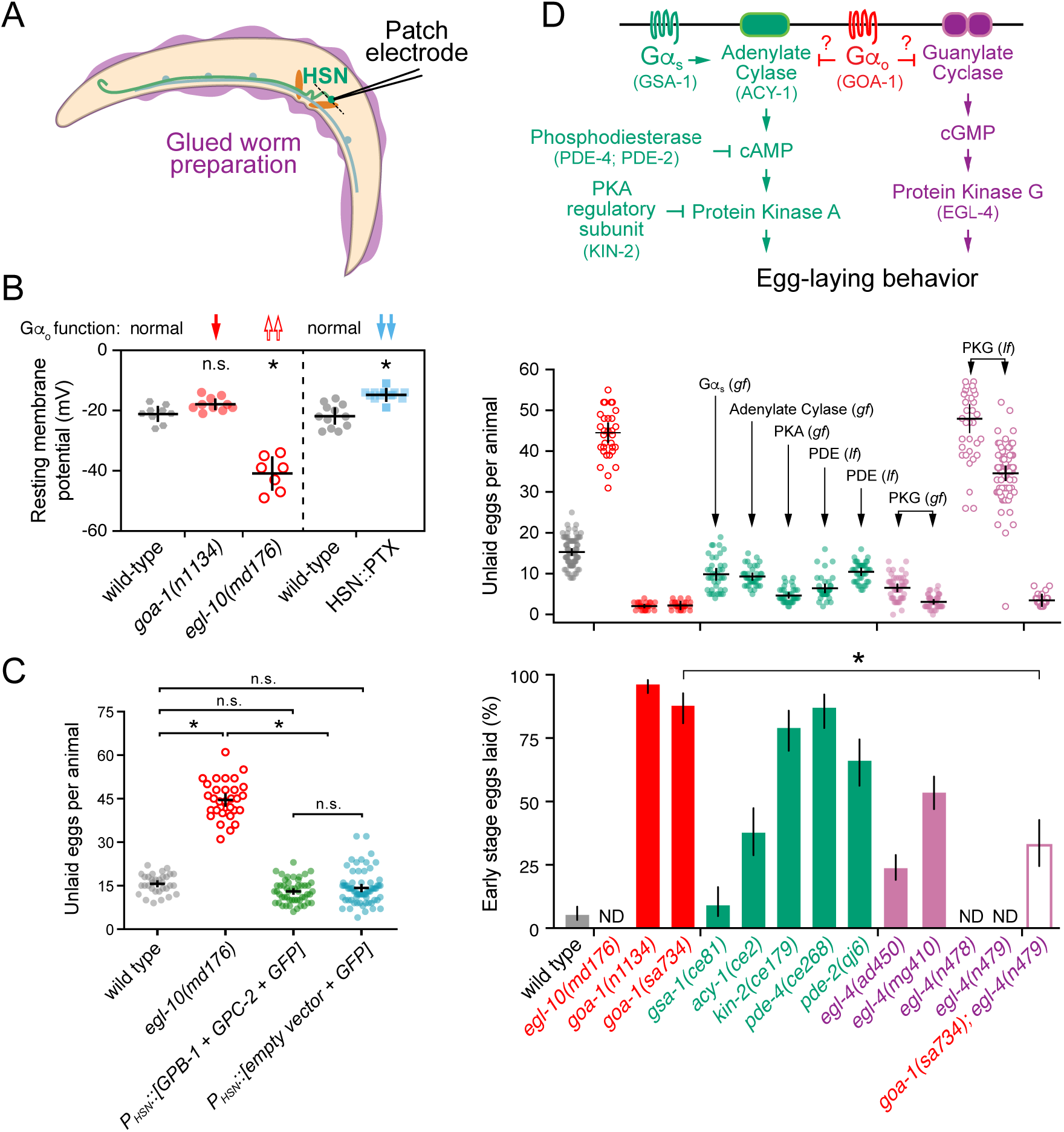
Gα_o_ depresses HSN resting membrane potential and may inhibit egg-laying behavior in parallel to βγ, cGMP, and cAMP signaling pathways. (**A**) Cartoon of the glued worm preparation used for patch clamp electrophysiology of HSN. (**B**) Scatter plots show resting membrane potential of wild-type control animals (grey circles), *goa-1(n1134)* loss-of-function mutants (red filled circles), *egl-10(md176)* null mutants (red open circles), and in transgenic animals expressing Pertussis Toxin in HSN (blue filled squares). Error bars indicate 95% confidence intervals for the mean. Asterisks indicate p<0.0001 (One-Way ANOVA with Bonferroni correction for multiple comparisons). N≥7 animals recorded per genotype. (**C**) Scatter plots show average number of eggs retained by wild-type animals (gray filled circles), *egl-10(md176)* null mutants (orange open circles), and in transgenic animals expressing either Gβ (GPB-2) and Gγ (GPC-1) subunits (green filled circles) or nothing (blue filled circles) in HSN from the *tph-1* gene promoter along with GFP. Error bars indicate means with 95% confidence intervals. Asterisk indicates p<0.0001; n.s. indicates p>0.05 (One-Way ANOVA with Bonferroni correction for multiple corrections). (**D**) Top, cartoon of how Gα_o_ signaling might interact with cAMP and cGMP signaling pathways. Gene names for *C. elegans* orthologs tested here are indicated in parentheses. Middle, scatterplots show average number of eggs retained by wildtype (grey), *egl-10(md176)* (red open circles), *goa-1(n1134)* and *goa-1(sa734)* Gα_o_ loss of function (red filled circles) mutants, and in animals with altered cAMP effector signaling (green): *gsa-1(ce81)* Gαs gain-of-function, *acy-1(ce2)* Adenylate Cyclase gain-of-function, *kin-2(ce179)* Protein Kinase A (PKA) inhibitory regulatory subunit loss-of-function, *pde-4(ce268)* Phosphodiesterase (PDE) loss-of-function, *pde-2(qj6)* Phosphodiesterase (PDE) null; altered cGMP effector signaling (pink): *egl-4(ad805)* and *egl-4(mg410)* Protein Kinase G (PKG) gain-of-function (pink filled circles), *egl-4(n478)* and *egl-4(n479)* loss-of-function mutants (pink open circles). Mean totals below ~10 eggs indicate hyperactive egg laying while totals above ~20 eggs indicate egg-laying behavior defects. Bottom, bar graphs indicate percent of embryos laid at early stages of development. Animals laying ≥50% embryos at early stages are considered hyperactive. Error bars indicate ± 95% confidence intervals for the mean proportion. N.D. indicates the stages of eggs laid was not determined because those mutants are egg-laying defective (Egl). Asterisk indicates highlighted significant differences (p≤0.0001; Fisher Exact Test with Bonferroni correction for multiple comparisons).

### Inhibition of egg laying by Gα_o_ is not replicated by elevated βγ expression

Receptor activation of Gα_i/o_ heterotrimers releases βγ subunits which have previously been shown to bind to activate specific K^+^ channels and inhibit Ca^2+^ channels (Reuveny et al., 1994; Herlitze et al., 1996). RNAi-mediated knockdown of the GPB-1 β subunit in HSN reduces egg-laying behavior (Esposito et al., 2007). An increase in free βγ subunits in Gα_o_ mutants might explain their hyperactive egg-laying behavior phenotypes. To test if βγ over-expression in HSN would increase egg laying, we transgenically overexpressed the sole non-RGS Gβ subunit, GPB-1 (Jansen et al., 1999), and the broadly expressed Gγ subunit, GPC-2 (Yamada et al., 2009), under the *tph-1* promoter along with GFP. βγ over-expression in HSN did not cause any significant differences in steady-state egg accumulation (Figure 5C). The number of eggs stored *in-utero* in these animals (13.0 ± 1.1 eggs) was comparable to wild-type animals (15.7 ± 1.2 eggs) and less than *egl-10(md176)* mutant animals (44.5 ± 2.3 eggs). These results suggest that Gα_o_ signals to inhibit HSN activity in a distinct manner from simple titration or release of βγ subunits.

### Egg-laying behavior is dysregulated in cAMP and cGMP signaling mutants

Gα_o_ is in the Gα_i/o_ class of G proteins that bind to and inhibit nucleotide cyclases, which function to reduce cAMP and cGMP levels and their subsequent activation of protein kinases (Kobayashi et al., 1990; Zhang and Pratt, 1996; Matsubara, 2002; Ghil et al., 2006). The hyperactive egg-laying behavior phenotypes of animals with reduced Gα_o_ function could be explained by a failure to properly terminate cAMP and/or cGMP signaling (Figure 5D, top). To test this, we analyzed egg-laying behavior in strains bearing mutations that increase Gαs and cAMP effector signaling (Reynolds et al., 2005; Schade et al., 2005; Charlie et al., 2006a; Charlie et al., 2006b). Animals carrying *gsa-1(ce81)* gain-of-function mutations in Gαs predicted to increase signaling accumulate fewer eggs compared to wild-type animals (Figure 5D, middle). Because a reduction in steady-state egg accumulation could be caused by indirect effects of these mutations on egg production or brood size, we also examined the developmental age of embryos laid. As previously reported (Bany et al., 2003), loss of inhibitory Gα_o_ signaling causes embryos to be laid precociously, before they reach the 8-cell stage (Figure 5D, bottom). Gαs gain-of-function mutant animals do not show a corresponding increase in early-stage embryos that are laid, suggesting the reduction in egg accumulation observed in Gαs gain-of-function mutant animals is indirect. In contrast, gain-of-function *acy-1* Adenylate Cyclase mutations or loss-of-function *pde-4* phosphodiesterase mutations, both predicted to result in increased cAMP signaling, cause animals to accumulate fewer eggs and lay them at earlier stages (Figure 5D). Similarly, *kin-2* mutations that disrupt an inhibitory regulatory subunit of Protein Kinase A showed a modest but significant hyperactive egg-laying phenotype. Together, these results indicate that Gαs, cAMP, and Protein Kinase A signaling promote egg-laying behavior. Because these mutant phenotypes were not nearly as strong as animals carrying Gα_o_ mutants, Gα_o_ likely signals to inhibit egg laying via other effectors.

Extensive work has shown that the cGMP-dependent Protein Kinase G, EGL-4, regulates egg laying in *C. elegans* (Fujiwara et al., 2002; L’Etoile et al., 2002; Raizen et al., 2006; Hao et al., 2011). Mutations which increase EGL-4 activity cause hyperactive egg laying and release of early stage embryos, while elimination of EGL-4 signaling causes egg-laying defects and significant egg accumulation (Figure 5D, middle). To determine whether Gα_o_ and Protein Kinase G regulate egg laying in a shared pathway, we performed a genetic epistasis experiment. *goa-1(sa734)*; *egl-4(n479)* double null mutants accumulate very few eggs (Figure 5D, middle), resembling the *goa-1(sa734)* null mutant. However, the low brood size of the *goa-1(sa734)* mutant could prevent accurate measurement of these animal’s egg-laying defects. To address this, we measured the stage of eggs laid as indicated above. Loss of the EGL-4 Protein Kinase G strongly and significantly suppressed the hyperactive egg-laying behavior of Gα_o_ null mutants (Figure 5D, bottom). *goa-1(sa734)*; *egl-4(n479)* mutants laid 33% of their embryos at early stages compared to 88% for the *goa-1(sa734)* single mutant, an intermediate egg-laying phenotype. These results are consistent with cGMP and/or Protein Kinase G signaling acting downstream of Gα_o_ in the proper regulation of egg-laying behavior.

### Gα_o_ signals in the vulval muscles and uv1 neuroendocrine cells to inhibit egg laying

In addition to HSN, GOA-1 is expressed in all neurons of the reproductive circuit, the egg-laying vulval muscles, and the uv1 neuroendocrine cells (Mendel et al., 1995; Segalat et al., 1995; Jose et al., 2007), raising questions as to how Gα_o_ signaling functions in those other cells to regulate egg-laying behavior. Previous work has shown that GOA-1(Q205L) was sufficient to rescue the hyperactive egg-laying behavior of *goa-1(n1134)* mutants when expressed in the HSNs but had little effect when expressed in the VCs or vulval muscles (Tanis et al., 2008). Similarly, transgenic expression of Pertussis Toxin in the VCs or vulval muscles failed to modify egg-laying behavior strongly (Tanis et al., 2008). Previous work failing to identify a function for Gα_o_ in the vulval muscles used a modified *Nde*-box element from the *ceh-24* promoter that drives expression more efficiently in the vm1 muscle cells compared to the vm2 muscles innervated by the HSN and VC neurons (data not shown). We find that expression of Pertussis Toxin in both vm1 and vm2 vulval muscles from a larger region of the *ceh-24* gene promoter (Harfe and Fire, 1998; Ravi et al., 2018a) also failed to cause any significant changes in the steady-state egg accumulation (Figure 6A). Conversely, expression of the activated GOA-1(Q205L) mutant in the vulval muscles from the same promoter did cause a modest but significant egg-laying defect, with animals accumulating 24.1 ± 2.0 eggs compared to mCherry-expressing control transgenic animals (13.2±0.7 eggs). This egg-laying defect was significantly weaker than *egl-10(md176)* mutants (46.0 ± 3.3 eggs) or transgenic animals expressing GOA-1(Q205L) just in the HSNs (36.8 ± 3.8 eggs). We do not believe this modest egg-laying defect was caused by transgene expression outside of the vulval muscles, as blocking synaptic transmission via transgenic expression of Tetanus Toxin from the same *ceh-24* promoter showed no such egg-laying defect (Figure 6B). Collectively, these results show that Gα_o_ does not play a significant role in suppressing vulval muscle excitability under state-state conditions but that activated Gα_o_ can signal in these cells to induce a mild but significant inhibition of cell activity and egg-laying behavior.

**Figure 6.**
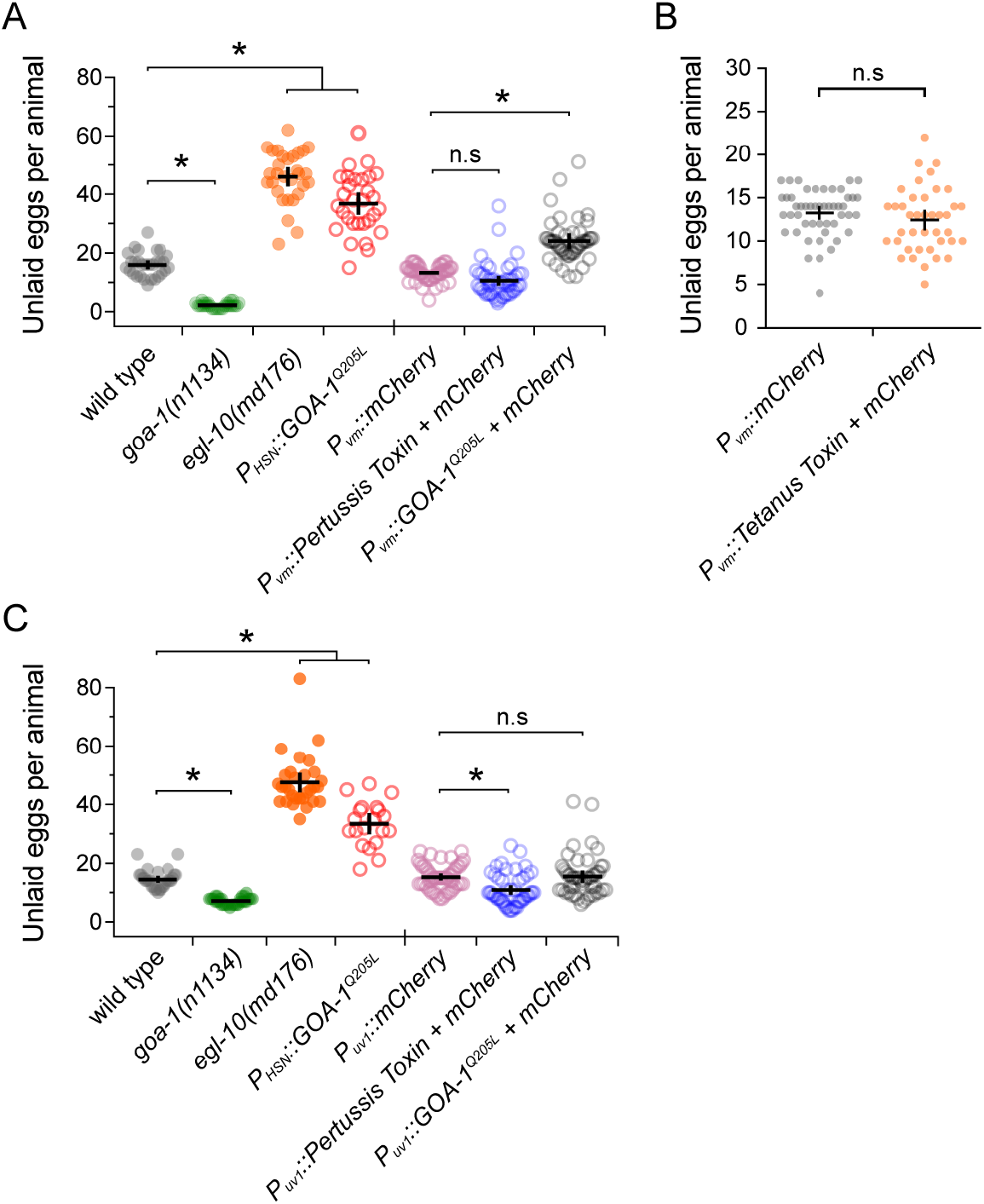
Altered Gα_o_ signaling in the vulval muscles and uv1 neuroendocrine cells causes only modest effects on egg laying. **(A)** Scatter plots show average number of eggs retained by wild-type (grey), *goa-1(n1134)* mutants (green), *egl-10(md176)* null mutant animals (orange) along with transgenic animals expressing GOA-1^Q205L^ in the HSNs (red open circles) compared to transgenic animals expressing mCherry only (pink open circles), Pertussis Toxin (blue open circles), or GOA-1^Q205L^ (black open circles) in the vulval muscles from the *ceh-24* gene promoter. (**B**) Scatter plots show average number of eggs retained in transgenic animals expressing mCherry only (gray) or Tetanus Toxin along with mCherry (orange) in the vulval muscles (vm) using the *ceh-24* gene promoter. Error bars indicate means with 95% confidence intervals. (**C**) Scatter plots show average number of eggs retained by wild-type (grey), *goa-1(n1134)* mutant (green), *egl-10(md176)* null mutant (orange) animals, and transgenic animals expressing GOA-1^Q205L^ in the HSNs (red) compared to transgenic animals expressing mCherry only (pink), Pertussis Toxin (blue), or GOA-1^Q205L^ (black open circles) in the uv1 neuroendocrine cells from the *tdc-1* gene promoter. Four or five independent extrachromosomal arrays were generated for each transgene in **(A-C)** and ~10 animals bearing each extrachromosomal array were analyzed. Error bars indicate 95% confidence intervals for the mean. Asterisks indicate p<0.0001; n.s. indicates p>0.05 (One-Way ANOVA with Bonferroni’s correction for multiple comparisons or Student’s t test).

The uv1 cells express the neurotransmitter tyramine along with NLP-7 and FLP-11 neuropeptides which inhibit egg laying (Alkema et al., 2005; Collins et al., 2016; Banerjee et al., 2017). Based on the function of Gα_o_ signaling in inhibiting neurotransmitter release in neurons, we would expect that loss of Gα_o_ function in uv1 would enhance their excitability, promoting release of inhibitory tyramine and neuropeptides, causing a reduction of egg laying. Surprisingly, previous work has shown that transgenic expression of Pertussis Toxin in uv1 cells increased the frequency of early-stage eggs that are laid, similar to the blocking of neurotransmitter release by Tetanus Toxin (Jose et al., 2007). A caveat of these experiments is that the *ocr-2* gene promoter used for transgene expression in uv1 also expresses in the utse (uterine-seam) associated cells and head sensory neurons (Jose et al., 2007). To test whether Gα_o_ functions specifically in uv1 to regulate egg laying, we used the *tdc-1* gene promoter (Alkema et al., 2005) along with the *ocr-2* 3’ untranslated region (Jose et al., 2007) to drive expression more specifically in uv1. Transgenic expression of Pertussis Toxin in uv1 caused a mild but significant decrease in steady-state egg accumulation (10.9 ± 1.5 eggs) compared to mCherry-expressing control animals (15.3 ± 1.2 eggs), indicating Gα_o_ signaling facilitates uv1-dependent inhibition of egg-laying behavior (Figure 6C). We also tested how increased Gα_o_ signaling in uv1 affects egg laying. Transgenic expression of the activated GOA-1(Q205L) mutant in uv1 cells caused no quantitative differences in egg accumulation (15.5 ± 2.1 eggs) (Figure 6C). Together, these results show that Gα_o_ has a limited role in regulating egg-laying behavior in the vulval muscles or uv1 neuroendocrine cells, unlike the strong phenotypes observed when we genetically manipulate Gα_o_ function in HSN.

### NLP-7 inhibition of HSN activity and egg-laying requires the EGL-47 receptor and Gα_o_

Multiple neuropeptides and receptors have been identified to inhibit egg laying by signaling through Gα_o_-coupled receptors expressed on HSN (Figure 1A). Recent work has identified NLP-7 neuropeptides, synthesized in the VC neurons and uv1 neuroendocrine cells, as potential ligands for EGL-47 receptor signaling through Gα_o_ (Moresco and Koelle, 2004; Banerjee et al., 2017). Animals overexpressing the NLP-7 neuropeptide are egg-laying defective, accumulating 39.6 ± 3.4 eggs in the uterus (Figure 7A). To test how NLP-7 inhibits egg laying, we crossed NLP-7 over-expressing transgenes into *goa-1* mutant animals and evaluated their egg-laying behavior phenotypes. As shown in Figure 7A, the hypomorphic *goa-1(n1134)* loss-of-function mutant significantly suppressed these egg-laying defects, as previously shown (Banerjee et al., 2017). However, the suppression by *goa-1(n1134)* was incomplete; NLP-7 over-expressing, *goa-1(n1334)* double mutant animals retained more eggs than the *goa-1(n1134)* single mutant or even wild-type control animals (Figure 7A). To confirm whether Gα_o_ was required for NLP-7 inhibition, we tested the *goa-1(sa734)* null mutant and found it fully suppressed the egg-laying defects caused by NLP-7 over-expression. To confirm this epistatic relationship, we measured the stage of embryos laid by these animals. *goa-1(sa734)* null mutant animals over-expressing NLP-7 laid ~100% of their embryos at early stages, a level not significantly different *goa-1(sa734)* single mutant animals (Figure 7B). Together, these results show that NLP-7 neuropeptides cannot inhibit egg-laying behavior in the absence of Gα_o_ function.

**Figure 7.**
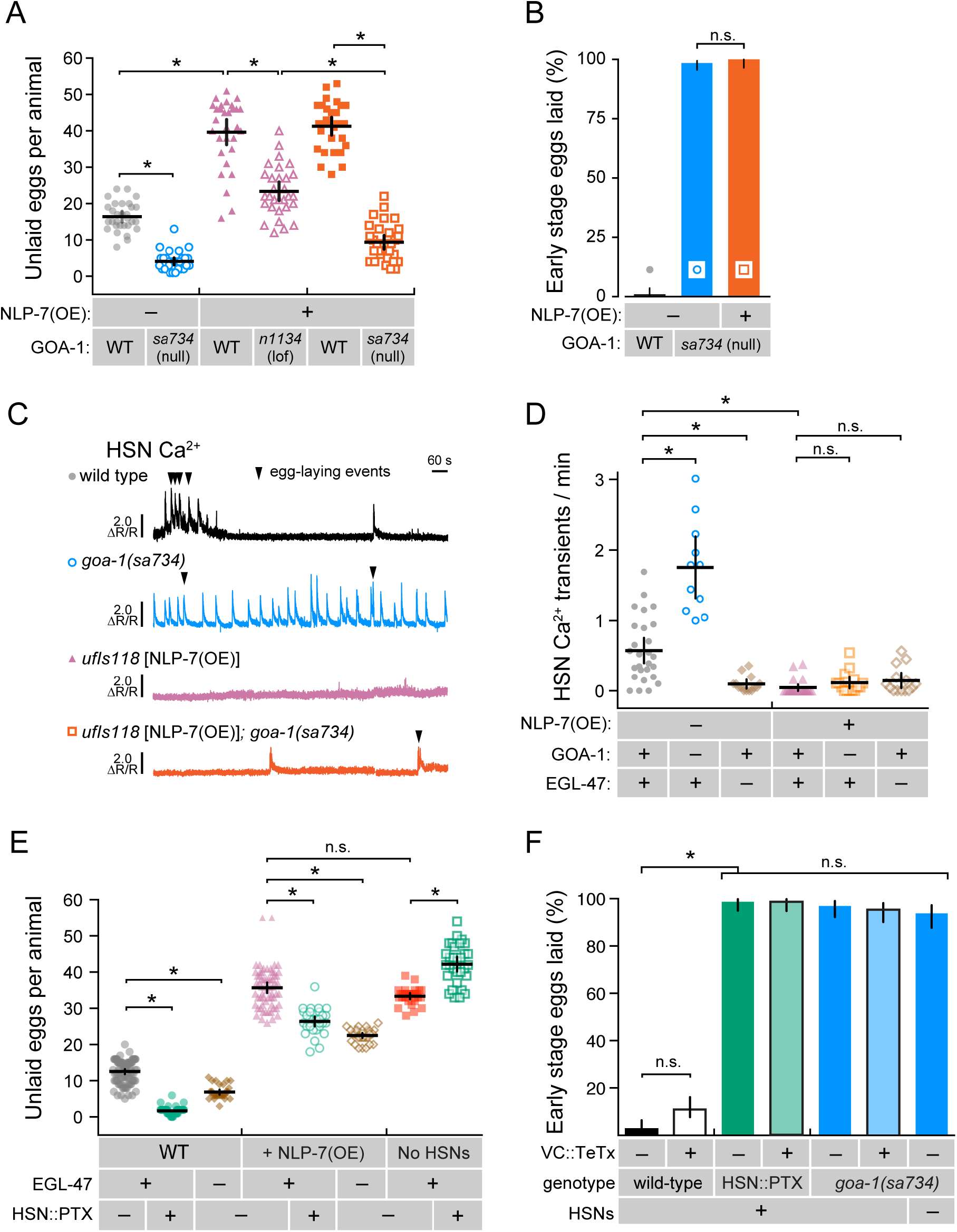
NLP-7 neuropeptides signal through EGL-47 and Gα_o_ to inhibit egg-laying outside of the HSNs. **(A)** NLP-7 signals through Gα_o_ to inhibit egg-laying behavior. Scatter plots show average number of eggs retained by wild-type (gray circles), *goa-1(sa734)* mutants (blue open circles), NLP-7 over-expressing (OE) transgenics in the wild-type (pink triangles or orange squares), and in NLP-7 over-expressing transgenics in the *goa-1(n1134)* (pink open triangles) and *goa-1(sa734)* null mutant background (orange open squares). Data in orange squares are from animals that also carry the *vsIs183* transgene used for HSN Ca^2+^ imaging. Mean totals below ~10 eggs indicate hyperactive egg laying while totals above ~20 eggs indicate egg-laying defective behavior defects. Error bars indicate 95% confidence intervals for the mean. Asterisks indicate p<0.0001 (One-Way ANOVA with Bonferroni correction for multiple comparisons). N≥30 animals for each strain. (**B**) Measure of early stage eggs laid by wild-type (black), *goa-1(sa734)* null mutants (blue), and *goa-1(sa734)* null mutants over-expressing NLP-7 neuropeptides (orange). Both *goa-1(sa734)* mutant strains also carry the *vsIs183* transgene used for HSN Ca^2+^ imaging. (**C**) NLP-7 over-expression silences HSN Ca^2+^ activity. Representative GCaMP5/mCherry ratio traces showing HSN Ca^2+^ activity in wild-type (black), *goa-1(sa734)* null mutants, and NLP-7 overexpressing transgenic animals in the wild-type (pink) and *goa-1(sa734)* null mutant backgrounds (orange). Arrowheads indicate egg-laying events. **(D)** Scatter plots show HSN Ca^2+^ peaks per minute measurements in wild-type (black), *goa-1(sa734)* null mutants, or *egl-47(ok677)* null mutant animals in the absence or presence of NLP-7 over-expression (OE). Error bars indicate 95% confidence intervals for the mean for N≥10 animals; n.s. indicates p>0.05 while asterisk indicates p≤0.0001 (one-way ANOVA with Bonferroni correction for multiple comparisons). **(E)** NLP-7 inhibition of egg laying is not exclusive to HSN silencing. Scatter plots show the average number of eggs retained by wild-type animals, transgenic animals expressing Pertussis Toxin in HSN, and *egl-47(ok677)* mutant animals in the absence or presence of NLP-7 over-expression (OE) or the absence of HSNs. Error bars indicate 95% confidence intervals for the mean. Asterisks indicate p<0.0001 (One-Way ANOVA with Bonferroni correction for multiple comparisons); N≥30 animals for each strain. (**F**) Measure of early stage eggs laid by wild-type or transgenic animals expressing Tetanus Toxin the VC neurons (black outlined boxes) in animals expressing Pertussis Toxin in the HSNs (green) or in *goa-1(sa734)* animals missing Gα_o_ (blue). Error bars indicate ± 95% confidence intervals for the mean proportion. Asterisk indicates highlighted significant differences (p≤0.0001) while n.s. indicates p>0.05 (Fisher Exact Test with Bonferroni correction for multiple comparisons).

Since the HSNs appear to be the principal sites of inhibitory Gα_o_ signaling, we tested how NLP-7 over-expression affects HSN Ca^2+^ activity. As expected, over-expression of NLP-7 strongly inhibited HSN Ca^2+^ activity (Figure 7C and 7D), consistent with the strong egg-laying defects of these animals. To our surprise, loss of Gα_o_ in these animals did now show high frequency HSN Ca^2+^ transient activity, despite showing the strong hyperactive egg-laying behavior of *goa-1(sa734)* single mutants (Figure 7C and 7D). This result shows that the hyperactive egg-laying behavior of Gα_o_ null mutants is unlinked to the increase in HSN Ca^2+^ activity we observe. Previous work showed NLP-7 inhibition of HSN activity and egg laying requires the EGL-47 receptor (Banerjee et al., 2017). To show directly whether NLP-7 inhibition of HSN activity requires the EGL-47 receptor, we recorded HSN Ca^2+^ activity in *egl-47(ok677)* null mutants. Unexpectedly, animals lacking EGL-47 had significantly fewer HSN Ca^2+^ transients than wild-type animals whether or not NLP-7 was over-expressed (Figure 7D). These results show that although NLP-7 signals to silence HSN Ca^2+^ activity, it does not require EGL-47 or Gα_o_ function to do so.

To confirm whether NLP-7 inhibition of egg laying requires EGL-47 and Gα_o_ function outside of HSN, we compared egg-laying behavior in NLP-7 transgenic animals where Gα_o_ and synaptic transmission were blocked in defined cells of the egg-laying circuit. As expected from our Ca^2+^ imaging experiments, NLP-7 over-expressing animals accumulated a similar number of eggs as *egl-1(n986dm)* animals lacking HSNs (35.7 ± 1.4 eggs vs. 33.4 ±0.9 eggs Figure 7E). Transgenic expression of Pertussis Toxin in HSNs or loss of the EGL-47 receptor weakly suppressed egg accumulation in NLP-7 over-expressing animals (Figure 7E). The suppression of NLP-7 egg-laying defects by Pertussis Toxin expression was specific, as *egl-1(n986dm)* animals developmentally lacking HSNs showed no comparable decrease in egg retention and showed a mild but significant increase (Figure 7E), possibly through co-expression of Pertussis Toxin from the transgene in the NSMs (Tanis et al., 2008). Thus, NLP-7 inhibits egg laying, in part, through an inhibitory G protein like Gα_o_ in HSN and the EGL-47 receptor. Because whole-animal, but not HSN-specific, elimination of Gα_o_ function completely suppressed NLP-7-induced egg-laying defects (Figure 7A), these data show that NLP-7 signals to inhibit egg laying through Gα_o_-coupled receptors on cells other than HSN. The cholinergic VC neurons also make extensive synapses onto the vulval muscles, suggesting they may act alongside the HSNs to promote vulval muscle contractility and egg laying (White et al., 1986; Waggoner et al., 1998). To test whether enhanced VC neurotransmitter release was driving the hyperactive egg-laying behavior of Gα_o_ signaling mutants, we transgenically expressed Tetanus Toxin in the VC neurons using a cell-specific promoter (Bany et al., 2003). Loss of VC synaptic transmission had no effect on egg-laying behavior by *goa-1(sa734)* null mutants or animals expressing Pertussis Toxin in HSN (Figure 7F). As previously reported (Mendel et al., 1995; Segalat et al., 1995), we find that loss of Gα_o_ also bypasses the regulation by of egg laying by the HSNs, as *goa-1(sa734)*; *egl-1(n986dm)* double mutants developmentally missing the HSNs still lay their eggs at early stages of development (Figure 7F). Together, these results show that NLP-7 neuropeptides and Gα_o_ signal to inhibit egg-laying behavior through cellular targets other than the VCs and HSNs.

## Discussion

Using a combination of genetic, imaging, physiological, and behavioral approaches, we found that the conserved G protein, Gα_o_, signals to depress activity in the *C. elegans* egg-laying circuit, preventing entry into the active behavior state. Neuropeptides, released in response to aversive external sensory input and feedback of successful egg release, activate inhibitory receptors and Gα_o_ which signal to reduce cell electrical excitability and neurotransmitter release. Without inhibitory Gα_o_ signaling, the presynaptic HSN command neurons remain electrically excited, rendering them resistant to inhibitory sensory input and feedback from the stretch-dependent homeostat. As a result, animals lacking Gα_o_ enter the egg-laying active state twice as frequently as wild-type animals and in environments that would normally be unfavorable for egg laying. Thus, inhibitory Gα_o_ signaling allows for a two-state pattern of circuit activity that can be properly gated by external and internal sensory input.

Our results inform our understanding of how G protein signaling modulates the stretch-dependent homeostat that governs egg-laying behavior. Even in ‘optimal’ environmental conditions including abundant food, egg laying in wild-type animals typically begins ~6 hours after the L4-adult molt upon the accumulation of 5-8 eggs in the uterus (Ravi et al., 2018a). Loss of inhibitory Gα_o_ signaling causes precocious egg laying, with eggs being released soon after being deposited into the uterus. Animals lacking Gα_o_ function in HSN show tonic Ca^2+^ transient activity even after chemical sterilization, conditions that normally inhibit HSN and vulval muscle Ca^2+^ activity in wild-type animals. Genetic perturbations that increase inhibitory Gα_o_ signaling delay egg laying until ~18 hours after the molt, like animals lacking HSNs altogether. Egg laying in animals with reduced or eliminated HSN function resumes when the accumulation of eggs activates the circuit via the stretch-dependent homeostat (Collins et al., 2016; Ravi et al., 2018a). Wild-type animals acutely exposed to aversive sensory conditions show a rapid inhibition of HSN activity and egg laying (Sawin, 1996; Zhang et al., 2008; Fenk and de Bono, 2015), but they continue to make and accumulate eggs in the uterus. Egg laying resumes upon removal of the inhibitory sensory stimulus or a return to food (Sawin, 1996; Dong et al., 2000). Feedback of egg accumulation and release has long-term consequences on female reproductive physiology. The cGMP-dependent Protein Kinase G ortholog, EGL-4, regulates the expression of a novel secreted protein in the uterine epithelium whose levels correlate with egg-laying rate (Hao et al., 2011). Feedback of egg release regulates *tph-1* gene expression that increases serotonin biosynthesis in the HSNs, alters sensory responses to male pheromone, and germline precursor cell proliferation (Aprison and Ruvinsky, 2019b, a). Because the primary function of all adult *C. elegans* behaviors is survival and reproduction, we hypothesize that most or all salient sensory signals ultimately converge to modulate egg-laying circuit activity and animal fecundity.

Our work shows that Gα_o_ also signals in cells other than HSN to inhibit egg-laying circuit activity and behavior. GOA-1 is likely expressed in all electrically excitable cells including all cells in the egg-laying circuit (Mendel et al., 1995; Segalat et al., 1995; Jose et al., 2007). Over-expression of NLP-7 neuropeptides strongly inhibits HSN Ca^2+^ activity and egg-laying behavior. While complete loss of Gα_o_ suppresses egg-laying defects caused by NLP-7 overexpression, HSN Ca^2+^ activity remains largely inhibited. This suggests NLP-7 signals through Gα_o_ to inhibit egg laying in cells other than HSN. Consistent with this result, *goa-1* null mutants remain strongly hyperactive for egg laying even when HSN or VC synaptic transmission is blocked. Where else does Gα_o_ function to inhibit egg laying? While Gα_o_ is expressed in the vulval muscles, transgenic expression of Pertussis Toxin or a GTP-locked form of Gα_o_ causes only modest egg-laying defects. Extensive work has shown that Gα_o_ signals to inhibit release of acetylcholine from motor neurons during locomotion (Miller et al., 1999; Nurrish et al., 1999). EM reconstruction that revealed the *C. elegans* synaptic wiring diagram shows that the VA7 and VB6 cholinergic motor neurons that drive body wall muscle contraction for locomotion also synapse onto the vm1 vulval muscles (White et al., 1986). Recent work employing a new fluorescent reporter of acetylcholine shows rhythmic activity in the vm1 muscles (Borden et al., 2020), and this vm1 activity is precisely where we observe weak vulval muscle ‘twitch’ Ca^2+^ transients (Collins et al., 2016; Brewer et al., 2019). Mutations that increase Gαs and cAMP signaling increase acetylcholine release from motor neurons and promote locomotion (Reynolds et al., 2005; Schade et al., 2005; Charlie et al., 2006a; Charlie et al., 2006b), possibly explaining why these mutations also increase egg laying. We predict that cholinergic neurons like VA7 and VB6, or their command interneurons, express neuropeptide and neurotransmitter receptors that signal through Gα_o_ to depress acetylcholine release and excitation of the vulval muscles. Identifying where such receptors are expressed and how they signal will help explain how each cell in the circuit functions to drive discrete steps in egg laying (Brewer et al., 2019; Fernandez et al., 2020).

Our work supports a model where Gα_o_ inhibits synaptic transmission via modulation of presynaptic ion channels. Genetic studies in *C. elegans* have identified the IRK inward rectifying K^+^ channels, NCA Na^+^ leak channels, and CCA-1 T-type voltage-gated Ca^2+^ channels as potential targets of Gα_o_ signaling (Yeh et al., 2008; Emtage et al., 2012; Topalidou et al., 2017a; Zang et al., 2017). Inward rectifying GIRK K^+^ channels are activated by release of βγsubunits (Hille, 1994). Previous work has shown the IRK-1 K^+^ channel is expressed in HSN and is required for inhibition of egg laying by the Gα_o_-coupled EGL-6 neuropeptide receptor (Emtage et al., 2012). The egg-laying phenotypes of *irk-1* mutant animals are not as strong as *goa-1* mutants, and over-expression of βγ subunits in HSN causes little or no effect on egg laying. These results suggest Gα_o_ signals to inhibit HSN neurotransmitter release via additional mechanism(s). NALCN Na^+^ leak channels are also expressed in HSN, and gain-of-function mutations increase HSN Ca^2+^ activity and drive hyperactive egg-laying and locomotion behaviors (Yeh et al., 2008). NALCN channels are genetically downstream of both Gα_q_ and Gα_o_ in the regulation of dopamine signaling and locomotion, suggesting that NALCN channels could be direct targets for modulation (Lutas et al., 2016; Topalidou et al., 2017b). Physiological experiments in mammalian neurons support this model. NALCN currents are activated by neuropeptide signaling through Gα_q_ (Lu et al., 2009) and inhibited by dopamine and GABA signaling through Gα_i/o_ (Philippart and Khaliq, 2018). However, *C. elegans* knockout mutants in NALCN channel components do not show the strong behavior defects seen in Gα_q_ and Gα_o_ mutants, suggesting other channels act in parallel to NALCN to regulate HSN and circuit excitability. One candidate are TMC channels that drive a background Na^+^ leak conductance that promotes HSN and vulval muscle cell electrical excitability (Yue et al., 2018). TMC channels may also regulate cell electrical excitability as mechanosensors (Pan et al., 2018; Tang et al., 2020). Cl^-^ channels and transporter proteins also regulate HSN activity and egg-laying behavior. CLH-3 is a swelling and hyperpolarization-activated, inwardly rectifying chloride channel that inhibits HSN activity (Branicky et al., 2014). Dynamic expression of Cl^-^ transporter proteins in the HSNs as animals mature into egg-laying adults regulates the Cl^-^ reversal potential (Tanis et al., 2009; Bellemer et al., 2011; Han et al., 2015). Loss of KCC-2 or ABTS-1 transporters flips the reduced egg laying behavior of *egl-47(dm)* mutants into hyperactive egg laying. Modulation of Cl^-^ transporter expression or activity by G protein signaling could drive the shifts in cell excitability we observe between the inactive and active egg-laying behavior states. The modulation of antagonistic cation and anion channels likely determines the Ca^2+^ dependent spiking probability of cells in the egg-laying circuit. HSN expresses L-type, P/Q-type, and T-type voltage-gated Ca^2+^ channels (Mathews et al., 2003; Zang et al., 2017), and each contributes to a normal serotonin response and egg-laying behavior (Schafer and Kenyon, 1995; Lee et al., 1997; Kwok et al., 2006). The modest but significant changes in HSN resting membrane potential we observe in Gα_o_ signaling mutants are expected to alter the probability of eliciting Ca^2+^ spiking activity. Ca^2+^-dependent action potentials have been found in *C. elegans* neurons and muscles (Gao and Zhen, 2011; Liu et al., 2011; Liu et al., 2018) and is regulated by both L-type and T-type Ca^2+^ channels. T-type channels like CCA-1 can contribute to a ‘window current’ where the channel can pass current at depolarized potentials that are insufficient to trigger channel inactivation (Williams et al., 1997; Crunelli et al., 2005; Zang et al., 2017). Activation of these window currents might allow neurons like HSN to shift from spontaneous tonic firing to high frequency Ca^2+^ bursting. Future work leveraging the powerful molecular tools uniquely available in *C. elegans* and the egg-laying circuit along with direct physiological measurements of membrane potential will allow mechanistic insight into how neuromodulators like serotonin and neuropeptides signal through effectors to shape patterns of circuit activity that underlie distinct behavior states.

## Materials and Methods

### Nematode Culture and Developmental Staging

*Caenorhabditis elegans* hermaphrodites were maintained at 20°C on Nematode Growth Medium (NGM) agar plates with *E. coli* OP50 as a source of food as described (Brenner, 1974). For assays involving young adults, animals were age-matched based on the timing of completion of the L4 larval molt. All assays involving adult animals were performed using age-matched adult hermaphrodites 20-40 hours past the late L4 stage. Table 2 lists all strains used in this study and their genotypes.

**Table 2.** Strain names and genotypes for all animals used in this study (behavior assays and calcium imaging)

| Strain | Feature | Genotype | Reference |
| --- | --- | --- | --- |
| N2 | Bristol wild-type strain | wild type | (Brenner, 1974) |
| DA521 | EGL-4 (Protein Kinase G) gain-of-function mutant. Hyperactive egg laying, sluggish locomotion | *egl-4(ad805) IV* | (Raizen et al., 2006) |
| DG1856 | GOA-1 ($G\alpha_o$) null mutant, hyperactive egg laying | *goa-1(sa734) I* | (Robatzek and Thomas, 2000) |
| IZ1236 | NLP-7 overexpression | *ufls118* | (Banerjee et al., 2017) |
| KG421 | GSA-1 ($G\alpha_s$) gain-of-function mutant. Hyperactive locomotion | *gsa-1(ce81) I* | (Schade et al., 2005) |
| KG518 | ACY-1 (Adenylate Cyclase) gain-of-function mutant. Hyperactive locomotion | *acy-1(ce2) III* | (Schade et al., 2005) |
| KG532 | KIN-2 (Protein Kinase A inhibitory regulatory subunit) loss-of-function mutant. Hypersensitive to stimuli | *kin-2(ce179) X* | (Schade et al., 2005) |
| KG744 | PDE-4 (phosphodiesterase) loss-of-function mutant, Hyperactive locomotion | *pde-4(ce268) II* | (Charlie et al., 2006a) |
| LX849 | HSN and NSM activated GOA-1(Q205L), egg laying defective | *vsIs49; lin-15(n765ts) X* | (Tanis et al., 2008) |
| LX850 | HSN and NSM Pertussis Toxin, hyperactive egg laying | *vsIs50; lin-15(n765ts) X* | (Tanis et al., 2008) |
| LX1832 | Strain for transgene production, blue-light insensitive, multi-vulva at 20°C in the absence of *lin-15(+)* rescue transgene. | *lite-1(ce314) lin-15(n765ts) X* | (Gurel et al., 2012) |
| LX1918 | vulval muscle GCaMP5, mCherry | *vsIs164 lite-1(ce314) lin-15(n765ts) X* | (Collins et al., 2016) |
| LX1919 | Vulval muscle GCaMP5, mCherry | *vsIs165; lite-1(ce314) lin-15(n765ts) X* | (Collins et al., 2016) |
| LX2004 | HSN GCaMP5, mCherry | *vsIs183 lite-1(ce314) lin-15(n765ts) X* | (Collins et al., 2016) |
| LX2007 | HSN GCaMP5, mCherry | *vsIs186 lite-1(ce314) lin-15(n765ts) X* | (Collins et al., 2016) |
| MIA26 | no HSNs | *egl-1(n986dm) V* | (Garcia and Collins, 2019) |
| MIA36 | EGL-4 (Protein Kinase G) gain-of-function mutant. Hyperactive egg laying, sluggish locomotion | *egl-4(mg410) IV* | (Hao et al., 2011) |
| MIA123 | no HSNs with *lin-15* marked transgenes | *egl-1(n986dm) V; lite-1(ce314) lin-15(n765ts) X* | this study |
| MIA144 | VC Tetanus Toxin | *keyIs33; lite-1(ce314) lin-15(n765ts) X* | this study |
| MIA210 | GOA-1 ($G\alpha_o$) reduced-function mutant; HSN GCaMP5, mCherry | *goa-1(n1134) I; vsIs183 lite-1(ce314) lin-15(n765ts) X* | this study |
| MIA214 | GOA-1 ($G\alpha_o$) reduced-function mutant; vulval muscle GCaMP5, mCherry | *goa-1(n1134) I; vsIs164 lite-1(ce314) lin-15(n765ts) X* | this study |
| MIA216 | Increased $G\alpha_o$ signaling; HSN GCaMP5, mCherry | *egl-10(md176) V; vsIs183 lite-1(ce314) lin-15(n765ts) X* | this study |
| MIA218 | HSN and NSM Pertussis Toxin in blue-light insensitive, *lin-15* multi-vulva background | *vsIs50 lite-1(ce314) lin-15(n765ts) X* | this study |
| MIA227 | HSN and NSM Pertussis Toxin; HSN GCaMP5, mCherry | *vsIs186; vsIs50 lite-1(ce314) lin-15(n765ts) X* | this study |
| MIA245 | HSN and NSM Pertussis Toxin; vulval muscle GCaMP5, mCherry | *vsIs50 X vsIs165; lite-1(ce314) lin-15(n765ts) X* | this study |
| MIA256 | Vulval muscle mCherry (*ceh-24* promoter) | *keyEx45; lite-1(ce314) lin-15(n765ts) X* | this study |
| MIA257 | Vulval muscle mCherry + Pertussis Toxin (*ceh-24* promoter) | *keyEx46; lite-1(ce314) lin-15(n765ts) X* | this study |
| MIA258 | Vulval muscle mCherry + Activated GOA-1(Q205L) (*ceh-24* promoter) | *keyEx47; lite-1(ce314) lin-15(n765ts) X* | this study |
| MIA259 | uv1 cells mCherry (*tdc-1* promoter, *ocr-2* 3' UTR) | *keyEx48; lite-1(ce314) lin-15(n765ts) X* | this study |
| MIA260 | uv1 cells mCherry + Pertussis Toxin (*tdc-1* promoter, *ocr-2* 3' UTR) | *keyEx49; lite-1(ce314) lin-15(n765ts) X* | this study |
| MIA261 | uv1 cells mCherry + Activated GOA-1(Q205L) (*tdc-1* promoter, *ocr-2* 3' UTR) | *keyEx50; lite-1(ce314) lin-15(n765ts) X* | this study |
| MIA262 | Vulval muscle mCherry + Tetanus toxin (*ceh-24* promoter) | *keyEx51; lite-1(ce314) lin-15(n765ts) X* | this study |
| MIA263 | GOA-1 ($G\alpha_o$) null mutant; HSN GCaMP5, mCherry | *goa-1(sa734) I; vsIs183 lite-1(ce314) lin-15(n765ts) X* | this study |
| MIA277 | Increased $G\alpha_o$ signaling in HSN; HSN GCaMP5, mCherry | *vsIs49/+, +/vsIs183 lite-1(ce314) lin-15(n765ts) X (trans-heterozygote)* | this study |
| MIA278 | HSN/NSM *gpb-1* and *gpc-2* overexpression + GFP (*tph-1* promoter) | *keyEx52; lite-1(ce314) lin-15(n765ts) X* | this study |
| MIA279 | HSN/NSM empty + GFP (*tph-1* promoter) | *keyEx53; lite-1(ce314) lin-15(n765ts) X* | this study |
| MIA282 | PDE-2 (phosphodiesterase) loss-of-function mutant | *pde-2(qj6)* | (Fujiwara et al., 2015) |
| MIA291 | Increased $G\alpha_o$ signaling in HSN; vulval muscle GCaMP5, mCherry | *vsIs49; vsIs165 lite-1(ce314) lin-15(n765ts) X* | this study |
| MIA293 | NLP-7 overexpression; HSN GCaMP5, mCherry | *ufls118; vsIs183 lite-1(ce314) lin-15(n765ts) X* | (Banerjee et al., 2017) & this study |
| MIA294 | NLP-7 overexpression; *goa-1(G$\alpha_o$)* null mutant; HSN GCaMP5, mCherry | *ufls118; goa-1(sa734) I; vsIs183 lite-1(ce314) lin-15(n765ts) X* | (Banerjee et al., 2017) & this study |
| MIA295 | GOA-1 ($G\alpha_o$) mutant, vulval muscles GCaMP5, mCherry | *goa-1(sa734) I; vsIs164 lite-1(ce314) lin-15(n765ts) X* | this study |
| MIA331 | GOA-1 ($G\alpha_o$) null mutant; EGL-4 (Protein Kinase G) null mutant | *goa-1(sa734)I; egl-4(n479)IV* | this study |
| MIA345 | GOA-1 ($G\alpha_o$) null mutant; no HSNs | *goa-1(sa734) I; egl-1(n986dm) V* | this study |
| MIA359 | EGL-47 null mutant; HSN GCaMP5, mCherry | *egl-47(ok677) V; vsIs183 lite-1(ce314) lin-15(n765ts) X* | this study |
| MIA360 | NLP-7 overexpression; HSN and NSM Pertussis Toxin; HSN GCaMP5, mCherry | *ufls118; vsIs50; vsIs186 lite-1(ce314) lin-15(n765ts) X* | this study |
| MIA366 | NLP-7 overexpression; EGL-47 null mutant; HSN GCaMP5, mCherry | *ufls118; egl-47(ok677) V; vsIs183 lite-1(ce314) lin-15(n765ts) X* | this study |
| MIA369 | no HSNs; HSN and NSM Pertussis Toxin; HSN GCaMP5, mCherry | *egl-1(n986dm) V; vsIs50; vsIs186 lite-1(ce314) lin-15 (n765ts) X* | this study |
| MIA370 | VC Tetanus Toxin; HSN and NSM Pertussis Toxin; HSN GCaMP5, mCherry | *keyIs33; vsIs186; vsIs50 keyIs33; lite-1(ce314) lin-15(n765ts) X* | this study |
| MIA371 | GOA-1 ($G\alpha_o$) null mutant; VC Tetanus Toxin | *goa-1(sa734) I; keyIs33; lite-1(ce314) lin-15(n765ts) X* | this study |
| MT1073 | EGL-4 (Protein Kinase G) loss-of-function mutant. Egg-laying defective, roaming locomotion | *egl-4(n478) IV* | (L'Etoile et al., 2002) |
| MT1074 | EGL-4 (Protein Kinase G) null mutant. Egg-laying defective, roaming locomotion | *egl-4(n479) IV* | (L'Etoile et al., 2002) |
| MT2426 | GOA-1 ($G\alpha_o$) reduced-function mutant, hyperactive egg laying | *goa-1(n1134) I* | (Segalat et al., 1995) |
| MT8504 | Increased $G\alpha_o$ signaling due to mutation in RGS protein, EGL-10 | *egl-10(md176) V* | (Koelle and Horvitz, 1996) |
| RB850 | EGL-47 null mutant | *egl-47(ok677) V* | (Moresco and Koelle, 2004; Consortium, 2012) |

### Plasmid and Strain Construction

Calcium reporter transgenes

#### HSN Ca^2+^

HSN Ca^2+^ activity was visualized using LX2004 *vsIs183* [*nlp-3::GCaMP5::nlp-3 3’UTR + nlp-3::mCherry::nlp-3 3’UTR + lin-15(+)*] *lite-1(ce314) lin-15(n765ts) X* strain expressing GCaMP5G and mCherry from the *nlp-3* promoter as previously described (Collins et al., 2016). To visualize HSN Ca^2+^ activity in Gα_o_ signaling mutants, we crossed LX2004 *vsIs183 lite-1(ce314) lin-15(n765ts) X* males with MT2426 *goa-1(n1134) I*, DG1856 *goa-1(sa734) I*, or MT8504 *egl-10(md176) V* hermaphrodites, and the fluorescent cross-progeny were allowed to self, generating MIA210 *goa-1(n1143) I*; *vsIs183 X lite-1(ce314) lin-15 (n765ts) X,* MIA263 *goa-1(sa734) I*; *vsIs183 X lite-1(ce314) lin-15 (n765ts) X*, and MIA216 *egl-10(md176) V; vsIs183 lite-1(ce314) lin-15(n765ts) X* strains, respectively. To visualize how NLP-7 signals to inhibit HSN Ca^2+^ activity and egg laying, we crossed IZ1236 *ufIs118* hermaphrodites (Banerjee et al., 2017) with LX2004 males. The fluorescent cross-progeny were selfed, generating MIA293 *ufIs118; vsIs183 lite-1(ce314) lin-15(n765ts) X.* To test the requirement for Gα_o_ signaling in NLP-7 inhibition, we crossed N2 males into MIA263 hermaphrodites, and the *goa-1(sa734)*/+ I; *vsIs183 lite-1(ce314) lin-15 (n765ts) X* heterozygous males obtained were then crossed with MIA293 hermaphrodites to generate MIA294 *ufIs118; goa-1(sa734) I; vsIs183 lite-1(ce314) lin-15(n765ts) X*. In order to test whether EGL-47 was a potential receptor for NLP-7, we crossed LX2004 males into RB850 *egl-47(ok677) V* hermaphrodites. The fluorescent cross-progeny were allowed to self and homozygous *egl-47(ok677)* mutant animals were identified by duplex PCR genotyping with the following oligos: CGTCACTTTTCCGTTGCTCTCTCATG, GTAGCGGGAAAGATGGCAAGAAGTCG, and TCGGTGAAACTCATTGTGCTCATTGTGC, creating MIA359 *egl-47(ok677) V; vsIs183 lite-1(ce314) lin-15(n765ts) X.* Wild-type N2 males were then crossed to MIA359 hermaphrodites, and the male *egl-47(ok677)/+ V; vsIs183 lite-1(ce314) lin15(n765ts) X* cross progeny were then crossed with MIA293 hermaphrodites. The cross-progeny were selfed and genotyped for the *egl-47(ok677) V* deletion mutant, generating MIA366 *ufIs118; egl-47(ok677) V; vsIs183 lite-1(ce314) lin-15(n765ts) X.*

During the course of crossing the *vsIs50* transgene expressing the catalytic subunit of Pertussis Toxin in the HSNs from the *tph-1* promoter (Tanis et al., 2008), we noticed repulsion between *vsIs183*, suggesting both were linked to the X chromosome. As such, LX850 *vsIs50 lin-15(n765ts) X* males were crossed with LX1832 *lite-1(ce314) lin-15(n765ts) X* hermaphrodites, the non-Muv progeny were selfed, and homozygous *lite-1(ce314)* non-Muv animals were kept, generating the strain MIA218 *vsIs50 lite-1(ce314) lin-15(n765ts) X.* MIA218 males were then crossed with LX2007 *vsIs186; lite-1(ce314) lin-15(n765ts) X*; the cross-progeny were selfed, generating MIA227 *vsIs186; vsIs50 lite-1(ce314) lin-15(n765ts) X*. To test whether Gα_o_ acts in HSN to mediate NLP-7 inhibition, we crossed wild-type N2 males into LX2007 *vsIs186; lite-1(ce314) lin-15 (n765ts) X* hermaphrodites, and the heterozygous *vsIs186*/+; *lite-1(ce314) lin-15 (n765ts) X* cross-progeny were mated with IZ1236 *ufIs118* hermaphrodites. The fluorescent cross-progeny were selfed, creating MIA358 *ufIs118; vsIs186 lite-1(ce314) lin-15(n765ts) X*. Wild-type N2 males were then crossed into MIA227 hermaphrodites to generate heterozygous *vsIs186*/+*; vsIs50 lite-1(ce314) lin-15(n765ts) X* males which were then crossed into MIA358 hermaphrodites. The fluorescent cross-progeny were selfed, generating MIA360 *ufIs118; vsIs50; vsIs186; lite-1(ce314) lin-15(n765ts) X*. Heterozygous *vsIs186*/+*; vsIs50 lite-1(ce314) lin-15(n765ts) X* males were also crossed into MIA123 *egl-1(n986dm)* V; *lite-1(ce314) lin-15(n765ts)* X hermaphrodites, the fluorescent cross-progeny were selfed, and Egl non-Muv animals were selected, generating MIA369 *egl-1(n986dm) V; vsIs50; vsIs186 lite-1(ce314) lin-15 (n765ts) X*. The presence of the *egl-1(n986dm)* allele was confirmed with snip-SNP PCR genotyping using oligonucleotides CTCTGTTCCAGCTCAAATTTCC and GTAGTTGAGGATCTCGCTTCGGC followed by NciI digestion.

In order to visualize HSN Ca^2+^ activity in transgenic animals expressing a constitutively active mutant GOA-1^Q205L^ protein that increases Gα_o_ signaling in the HSN neurons, LX2004 *vsIs183 lite-1(ce314) lin-15(n765ts) X* males were crossed with LX849 *vsIs49; lin-15(n765ts) X* hermaphrodites. We noted a repulsion between *vsIs183* and *vsIs49* transgenes integrated on X. As such, we selected a strain MIA277 with trans-heterozygous *vsIs49* and *vsIs183* transgenes (*lite-1(ce314) vsIs49 X / lite-1(ce314) vsIs183 X*) for Ca^2+^ imaging. The MIA277 strain was maintained by picking phenotypically egg-laying defective adult animals which show GCaMP5G/mCherry expression.

#### Vulval Muscle Ca^2+^

Vulval muscle Ca^2+^ activity was recorded in adult animals using LX1918 *vsIs164* [*unc-103e::GCaMP5::unc-54 3’UTR + unc-103e::mCherry::unc-54 3’UTR + lin-15(+)*] *lite-1(ce314) lin-15(n765ts) X* strain as described (Collins et al., 2016). To visualize vulval muscle activity in Gα_o_ signaling mutants, LX1918 males were crossed with MT2426 *goa-1(n1134)* I and DG1856 *goa-1(sa734) I* hermaphrodites, and the fluorescent cross-progeny were selfed, generating MIA214 *goa-1(n1134) I; vsIs164 lite-1(ce314) lin-15(n765ts) X* and MIA295 *goa-1(sa734) I*; *vsIs164 lite-1(ce314) lin-15(n765ts) X* strains, respectively. To visualize vulval muscle activity in transgenic animals expressing the catalytic subunit of Pertussis Toxin in the HSN neurons (Tanis et al., 2008), MIA218 *vsIs50 lite-1(ce314) lin-15(n765ts) X* males were crossed with LX1919 *vsIs165* [*unc-103e::GCaMP5::unc-54 3’UTR + unc-103e::mCherry::unc-54 3’UTR + lin-15(+)*]*; lite-1(ce314) lin-15(n765ts) X* hermaphrodites, and the cross progeny were selfed, generating MIA245 *vsIs50; vsIs165; lite-1(ce314) lin-15(n765ts) X*. To visualize vulval muscle activity in transgenic animals expressing a constitutively active mutant GOA-1^Q205L^ protein which increases Gα_o_ signaling in the HSN neurons (Tanis et al., 2008), LX849 *vsIs49; lin-15(n765ts) X* males were crossed with LX1919 *vsIs165; lite-1(ce314) lin-15(n765ts) X* hermaphrodites and the fluorescent cross-progeny were selfed, generating MIA291 *vsIs165; vsIs50 lite-1(ce314) lin-15(n765ts) X*.

Transgenes used to manipulate Gα_o_ signaling and synaptic transmission in specific cells of the egg-laying circuit

#### HSN neurons

To produce a HSN (and NSM)-specific GPB-1 expressing construct, the *gpb-1* cDNA fragment was amplified from pDEST-*gpb-1* (Yamada et al., 2009) using the following oligonucleotides: 5’-GAGGCTAGCGTAGAAAAAATGAGCGAACTTGACCAACTTCGA-3’ and 5’-GCGGGTACCTCATTAATTCCAGATCTTGAGGAACGAG-3’. The ~1 kb DNA fragment was digested with NheI/KpnI and ligated into pJT40A (Tanis et al., 2008) to generate pBR30. To produce an HSN (and NSM)-specific GPC-2 expressing construct, the *gpc-2* cDNA fragment was amplified from worm genomic DNA using the following forward and reverse oligonucleotides: 5’-GAGGCTAGCGTAGAAAAAATGGATAAATCTGACATGCAACGA-3’ and 5’-GCGGGTACCTTAGAGCATGCTGCACTTGCT-3’. The ~250 bp DNA fragment was digested with NheI/KpnI and ligated into pJT40A to generate pBR31. To co-overexpress the βγ G protein subunits in the HSN neurons, we injected pBR30 (50 ng/µl), pBR31 (50 ng/µl), and pJM60 [*ptph-1*::GFP] (80 ng/µl) (Moresco and Koelle, 2004) into the LX1832 *lite-1(ce314) lin-15(n765ts)* animals along with pLI5EK (50 ng/µl), generating five independent extrachromosomal transgenic lines which were used for behavioral assays. One representative transgenic strain, MIA278 [*keyEx52*; *lite-1(ce314) lin-15(n765ts)*], was kept. To generate a control strain for comparison in the egg-laying assays, we injected pJM66 [*ptph-1*::empty] (100 ng/µl) (Tanis et al., 2008) and pJM60 (80 ng/µl) into the LX1832 *lite-1(ce314) lin-15(n765ts)* animals along with pLI5EK (50 ng/µl) generating five independent extrachromosomal control transgenes which were used for behavioral assays. One representative transgenic strain, MIA279 [*keyEx53*; *lite-1(ce314) lin-15(n765ts)*], was kept. To test how loss of HSNs affected the hyperactive egg-laying behavior of *goa-1(sa734)* null mutants, we mated wild-type N2 males into MIA26 *egl-1(n986dm) V* hermaphrodites, and the heterozygous *egl-1(n986dm)*/+ male cross-progeny were then crossed into DG1856 *goa-1(sa734)* I hermaphrodites. Their cross-progeny were selfed, and the presence of the *egl-1(n986dm)* and *goa-1(sa734)* were confirmed by genotyping, creating MIA345 *goa-1(sa734)* I; *egl-1(n986dm)* V.

#### Vulval muscles

pJT40A *Ptph-1*::Pertussis Toxin (Tanis et al., 2008) was digested with NheI/KpnI and ligated into pBR3 (*Pceh-24*::mCherry) to generate pBR20. pBR20 [*Pceh-24*::Pertussis Toxin] (10 ng/µl) and pBR3 [*Pceh-24*::mCherry] (10 ng/µl) were injected into the LX1832 *lite-1(ce314) lin-15(n765ts)* animals along with pLI5EK (50 ng/µl) to generate five independent extrachromosomal transgenes which were used for behavioral assays. One representative transgenic strain, MIA257 [*keyEx46*; *lite-1(ce314) lin-15(n765ts)*], was kept. To produce vulval muscle-specific GOA-1(Q205L), the coding sequence of GOA-1(Q205L) was recovered from pJM70C (Tanis et al., 2008) after digestion with NheI/SacI and ligated into pKMC188 (*Punc-103e*::GFP; (Collins and Koelle, 2013)) generating pKMC268 (*Punc-103e::goa-1(Q205L)*). However, because the *unc-103e* promoter also directs expression in neurons that might indirectly regulate egg laying, GOA-1(Q205L) coding sequences were removed from pKMC268 by digesting with NheI/NcoI and ligated into pBR3 to generate pBR21. pBR21 [*Pceh-24*::GOA-1^Q205L^] (10 ng/µl) and pBR3 [*Pceh-24*::mCherry] (10 ng/µl) were injected into the LX1832 *lite-1(ce314) lin-15(n765ts)* animals along with pLI5EK (50 ng/µl) to generate five independent extrachromosomal transgenes which were used for behavior assays. One representative transgenic strain, MIA258 [*keyEx47*; *lite-1(ce314) lin-15(n765ts)*], was kept. To generate control strains for comparison in egg-laying assays, pBR3 [*Pceh-24*::mCherry] (20 ng/µl) was injected into the LX1832 *lite-1(ce314) lin-15(n765ts)* animals along with pLI5EK (50 ng/µl) to generate five independent extrachromosomal transgenes which were used for behavioral assays. One representative control transgenic strain, MIA256 [*keyEx45*; *lite-1(ce314) lin-15(n765ts)*], was kept. To produce a vulval muscle-specific Tetanus Toxin transgene, Tetanus Toxin coding sequences were amplified from pAJ49 (*Pocr-2*::Tetanus toxin) (Jose et al., 2007) using the following oligonucleotides: 5’-GAGGCTAGCGTAGAAAAAATGCCGATCACCATCAACAACTTC-3’ and 5’-GCGCAGGCGGCCGCTCAAGCGGTACGGTTGTACAGGTT-3’. The DNA fragment was digested with NheI/NotI and ligated into pBR6 to generate pBR27. To block any possible neurotransmitter release from the vulval muscles, pBR27 (10 ng/µl) and pBR3 (10 ng/µl) was injected into the LX1832 *lite-1(ce314) lin-15(n765ts)* animals along with pLI5EK (50 ng/µl) to generate five independent extrachromosomal transgenes which were used for behavior assays. One representative transgenic strain, MIA262 [*keyEx51*; *lite-1(ce314) lin-15(n765ts)*], was kept.

#### uv1 neuroendocrine cells

To generate a uv1 cell-specific Pertussis toxin transgene, pBR20 (*Pceh-24*::Pertussis toxin) was digested with NheI/NcoI and the coding sequences of Pertussis Toxin were then ligated into pAB5 (*Ptdc-1*::mCherry::*ocr-2* 3’UTR) to generate pBR25. pBR25 [*Ptdc-1*::Pertussis Toxin] (10 ng/µl) and pAB5 [*Ptdc-1*::mCherry] (5 ng/µl) were injected into LX1832 *lite-1(ce314) lin-15(n765ts)* animals along with pLI5EK (50 ng/µl) to generate five independent extrachromosomal transgenes which were used for behavioral assays. One representative transgenic strain, MIA260 [*keyEx49*; *lite-1(ce314) lin-15(n765ts)*], was kept. To generate a uv1 cell-specific GOA-1(Q205L) transgene, pKMC268 (*Punc-103e*::GOA-1(Q205L)) was digested with NheI/NcoI and the coding sequences of GOA-1(Q205L) were then ligated into pBR25 to generate pBR26. To increase Gα_o_ signaling in uv1 cells, we injected pBR26 [*Ptdc-1*::GOA-1^Q205L^] (10 ng/µl) and pAB5 [*Ptdc-1*::mCherry] (5 ng/µl) into the LX1832 *lite-1(ce314) lin-15(n765ts)* animals along with pLI5EK (50 ng/µl) to generate five independent extrachromosomal transgenes which were used for behavioral assays. One transgenic strain MIA261 [*keyEx50*; *lite-1(ce314) lin-15(n765ts)*] was kept. To generate a control strain for comparison in our egg-laying assays, pAB5 [*Ptdc-1*::mCherry] (15 ng/µl) was injected into the LX1832 *lite-1(ce314) lin-15(n765ts)* animals along with pLI5EK (50 ng/µl) to generate five independent extrachromosomal transgenes which were used for behavioral assays. One representative transgenic strain, MIA259 [*keyEx48*; *lite-1(ce314) lin-15(n765ts)*], was kept.

#### VC neurons

A ~1.4 kB DNA fragment encoding Tetanus Toxin was excised from pAJ49 (Jose et al., 2007) using AgeI/XhoI and ligated into a similarly digested pKMC145 *(Plin-11::GFP::unc-54 3’ UTR)* (Collins et al., 2016) to generate pKMC282 *(Plin-11::TeTx::unc-54 3’ UTR)*. pKMC282 (50 ng/µl) was injected along with pL15EK (50 ng/µl; (Clark et al., 1994) into LX1832 *lite-1(ce314) lin-15(n765ts)* X generating 4 independent transgenic lines of which one, MIA113 *keyEx32 [Plin-11::TeTx::unc-54 3’UTR + lin-15(+)],* was used for integration. *keyEx32* was integrated with UV/TMP generating three independent integrants (*keyIs32*, *keyIs33*, and *keyIs46*). Each transgenic line was backcrossed six times to LX1832 generating strains MIA144, MIA145, and MIA146. The behavioral characterization of these VC-silenced animals will be described elsewhere in more detail. To eliminate HSNs in these VC-silenced animals, wild-type N2 males were crossed into MIA26 *egl-1(n986dm) V* hermaphrodites, and the heterozygous *egl-1(n986dm)*/+ males were then crossed to MIA146 *keyIs46*; *lite-1(ce314) lin-15(n765ts) X* hermaphrodites, and the cross-progeny were then selfed, generating the homozygous strain MIA173 *keyIs46*; *egl-1(n986dm) V*; *lite-1(ce314) lin-15(n765ts) X*. To test whether synaptic transmission from the VCs was required for the hyperactive egg-laying behavior of Gα_o_ signaling mutants, wild-type N2 males were crossed to MIA227 *vsIs186; vsIs50 lite-1(ce314) lin-15(n765ts) X* hermaphrodites and the resulting heterozygous cross-progeny were then crossed MIA144 hermaphrodites, allowed to self, generating MIA370 *vsIs186; keyIs33; vsIs50 lite-1(ce314) lin-15(n765ts) X.* To test how complete loss of GOA-1 affected egg laying in VC-silenced animals, we crossed wild-type N2 males into DG1856 *goa-1(sa734)* I hermaphrodites, and the heterozygous *goa-1(sa734)*/+ cross-progeny were then mated with MIA144 hermaphrodites. The cross-progeny were allowed to self, generating MIA371 *goa-1(sa734) I; keyIs33; lite-1(ce314) lin-15(n765ts) X.* The presence of *goa-1(sa734)* was confirmed by Sanger sequencing.

### Fluorescence Imaging

#### Ratiometric Ca^2+^ Imaging

Ratiometric Ca^2+^ recordings were performed on freely behaving animals mounted between a glass coverslip and chunk of NGM agar, as previously described (Collins and Koelle, 2013; Li et al., 2013; Collins et al., 2016; Ravi et al., 2018b). Briefly, recordings were collected on an inverted Leica TCS SP5 confocal microscope using the 8 kHz resonant scanner at ~20 fps at 256×256 pixel resolution, 12-bit depth and ≥2X digital zoom using a 20x Apochromat objective (0.7 NA) with the pinhole opened to ~20 µm. GCaMP5G and mCherry fluorescence was excited using a 488 nm and 561 nm laser lines, respectively. Adult recordings were performed 24 hours after the late L4 stage. After staging, animals were allowed to adapt for ~30 min before imaging. During imaging, the stage and focus were adjusted manually to keep the relevant cell/pre-synapse in view and in focus.

Ratiometric analysis (GCaMP5:mCherry) for all Ca^2+^ recordings was performed after background subtraction using Volocity 6.3.1 as described (Ravi et al., 2018a). The egg-laying active state was operationally defined as the period one minute prior to the first egg-laying event and ending one minute after the last (in the case of a typical active phase where 3-4 eggs are laid in quick succession). However, in cases where two egg-laying events were apart by >60 s, peaks were considered to be in separate active phases and any transients observed between were considered to be from an inactive state. In animals where we observed no Ca^2+^ peaks during the entire recording, the total duration of the recording was considered as one, long, inter-transient interval. In animals where we observed a single Ca^2+^ transient, the duration from the start of the recording to the time of the Ca^2+^ transient and the time from the Ca^2+^ transient to the end of the recording were defined as inter-transient intervals. Animals overexpressing NLP-7 become highly egg-laying defective, and the embryonic expression of mCherry from the *nlp-3* promoter in accumulated eggs interfered with the standard image segmentation protocol based on the mCherry channel. For these animals, unambiguous Ca^2+^ transient-dependent changes in GCaMP5 fluorescence localized at the HSN cell body were scored manually, and the frequency of such events were calculated as the number of observed events per unit recording time, typically 10-20 minutes.

### Behavior Assays and Microscopy

#### Animal sterilization

Animals were sterilized using Floxuridine (FUDR) as described (Mitchell et al., 1979; Collins et al., 2016; Ravi et al., 2018a). Briefly, 100 µl of 10 mg/ml FUDR was applied to OP50 seeded NGM plates. Late L4 animals were then staged onto the FUDR plates and the treated adults were imaged ~24 hours later.

#### Egg laying assays

Unlaid eggs were quantitated as described (Chase et al., 2004). Staged adults were obtained by picking late L4 animals and culturing them for 30-40 hr at 20°C. The percentage of early-stage eggs laid were quantified as described (Koelle and Horvitz, 1996). 30 staged adults were placed on a thin lawn of OP50 bacteria on a nematode growth medium (NGM) agar plate (Brenner, 1974) and allowed to lay eggs for 30 min. This was repeated with new sets of staged animals until a total of at least 100 laid eggs were analyzed. Each egg was examined under a Leica M165FC stereomicroscope and categorized into the following categories: eggs which have 1 cell, 2 cell, 3-4 cell, 5-8 cell, and embryos with >8 cells. Eggs with eight cells or fewer were classified as “early stage.”

#### Long-term recording of egg-laying behavior

Egg-laying behavior was recorded at 4-5 frames per second from 24-hour adults after transfer to NGM plates seeded with a thin lawn of OP50 bacterial food using a Leica M165FC stereomicroscope and camera (Grasshopper 3, 4.1 Megapixel, USB3 CMOS camera, Point Grey Research). N2 wild-type and hyperactive egg-laying mutant strains (MT2426 and LX850) were recorded for 3 hours, and the egg-laying defective strains MT8504 and LX849 were recorded for 8-10 hours. Average duration of active and inactive states from these recordings were calculated as described (Waggoner et al., 1998; Waggoner et al., 2000).

### Electrophysiology

Electrophysiological recordings were carried out on an upright microscope (Olympus BX51WI) coupled with an EPC-10 amplifier and Patchmaster software (HEKA), as previously described (Yue et al., 2018; Zou et al., 2018). Briefly, day 2 adult worms were glued on the surface of Sylgard-coated coverslips using the cyanoacrylate-based glue (Gluture Topical Tissue Adhesive, Abbott Laboratories). A dorsolateral incision was made using a sharp glass pipette to expose the cell bodies of HSN neurons for recording. The bath solution contained (in mM) 145 NaCl, 2.5 KCl, 5 CaCl_2_, 1 MgCl_2_, and 20 glucose (325–335 mOsm, pH adjusted to 7.3). The pipette solution contained (in mM) 145 KCl, 5 MgCl_2_, 5 EGTA, 0.25 CaCl_2_, 10 HEPES, 10 glucose, 5 Na_2_ATP and 0.5 NaGTP (315–325 mOsm, pH adjusted to 7.2). The resting membrane potentials were tested with 0 pA holding under the Current Clamp model of whole-cell patch. The high Cl^-^ content of the intracellular solution used may contribute to the relatively depolarized membrane potentials we observed (Liu et al., 2018).

### Experimental Design and Statistical Analysis

Sample sizes for behavioral assays followed previous studies (Chase et al., 2004; Collins and Koelle, 2013; Collins et al., 2016). No explicit power analysis was performed before the study. Statistical analysis was performed using Prism v.6-8 (GraphPad). Ca^2+^ transient peak amplitudes and inter-transient intervals were pooled from multiple animals (typically ~10 animals per genotype/condition per experiment). No animals or data were excluded. Individual *p* values are indicated in each Figure legend, and all tests were corrected for multiple comparisons (Bonferroni for ANOVA and Fisher exact test; Dunn for Kruskal-Wallis).

## Acknowledgements

This work was funded by grants from the NIH (R01-NS086932) and NSF (IOS-1844657) to KMC. RJK 3^rd^ was supported by a University of Miami Maytag Fellowship. We thank Yuichi Iino for sharing plasmids. Strains used in this study have been provided to the *C. elegans* Genetics Center, which is funded by NIH Office of Research Infrastructure Programs (P40 OD010440). We thank Dr. Addys Bode Hernandez and Michael Scheetz for technical assistance. We thank Drs. Qiang Liu, James Baker, Julia Dallman, Laura Bianchi, Brock Grill, Peter Larsson, Stephen Roper, along with members of the Collins lab for helpful discussions and feedback on the manuscript.

---

**Video 1.** GCaMP5:mCherry ratio recording of HSN Ca^2+^ activity in a control, wild-type adult animal during an egg-laying active state. High Ca^2+^ is indicated in red while low calcium is in blue. Head is at left, tail is at right.

**Video 2.** GCaMP5:mCherry ratio recording of HSN Ca^2+^ activity in a *goa-1(n1134)* mutant adult animal during an egg-laying active state. High Ca^2+^ is indicated in red while low calcium is in blue. Head is at left, tail is at right.

**Video 3.** GCaMP5:mCherry ratio recording of HSN Ca^2+^ activity in a *goa-1(sa734)* null mutant adult animal during an egg-laying active state. High Ca^2+^ is indicated in red while low calcium is in blue. Head is at right, tail is at left.

**Video 4.** GCaMP5:mCherry ratio recording of HSN Ca^2+^ activity in a transgenic adult animal expressing Pertussis Toxin in the HSNs from the *tph-1* gene promoter during an egg-laying active state. High Ca^2+^ is indicated in red while low calcium is in blue. Head is at top, tail is at bottom.

**Video 5.** GCaMP5:mCherry ratio recording of vulval muscle Ca^2+^ activity in a control wild-type adult animal during an egg-laying active state. High Ca^2+^ is indicated in red while low calcium is in blue. Head is at right, tail is at top.

**Video 6.** GCaMP5:mCherry ratio recording of vulval muscle Ca^2+^ activity in a *goa-1(n1134)* mutant adult animal during an egg-laying active state. High Ca^2+^ is indicated in red while low calcium is in blue. Head is at right, tail is at left.

**Video 7.** GCaMP5:mCherry ratio recording of vulval muscle Ca^2+^ activity in a transgenic adult animal expressing Pertussis Toxin in the HSNs from the *tph-1* gene promoter during an egg-laying active state. High Ca^2+^ is indicated in red while low calcium is in blue. Head is at left, tail is at right.

